# A systematic approach for the purification of fluorophore-labelled proteins via anion exchange chromatography

**DOI:** 10.64898/2025.12.30.697019

**Authors:** Nicolas D. Wendler, Thorben Cordes

## Abstract

Förster resonance energy transfer (FRET), an optical distance ruler, provides the unique ability to monitor molecular interactions and conformational changes in single biomacromolecules and multi-subunit complexes. It has proven to be a powerful tool to study enzymatic reactions, membrane transport, protein folding, but also the properties of nucleic acids, proteins, molecular motors and many other biological systems and processes. A prerequisite for FRET is targeted (covalent) labelling of macromolecules with two distinct fluorophores. Here, we present a strategy for stochastic labeling of protein residues with the required donor and acceptor dyes via anion exchange chromatography. While this technique has been used before for this purpose, we provide a conceptual basis to systematically design a purification protocol for an arbitrary choice of fluorophores. By characterizing the interaction fluorophore-maleimides with the column material, we are able to select (and predict) which pairs of fluorophores allow successful purification of donor-acceptor-labelled protein with yields up to 98%. We demonstrate the capabilities of the method for bulk and single-molecule FRET assays of various bacterial substrate-binding proteins.

## Introduction

Advances in biophysical techniques[1–5] have steadily extended our understanding of fundamental mechanisms of biomolecular structure and function. While high-resolution structures obtained by X-ray crystallography[6,7], cryo-electron microscopy[8,9] and NMR[10–13] are still the cornerstone to unravel structure-function relationships, these techniques have limitations in resolving dynamic conformational changes, heterogeniety and (transient) interactions between biomolecules[14]. Förster Resonance Energy Transfer (FRET[15,16]), a non-radiative transfer of excitation energy between two optically addressable fluorophores has become an important technique to study exactly such processes in bulk and at the single-molecule level (smFRET[14,17]). FRET assays use the distance dependence of energy transfer on the 3-10 nm scale[18] between a donor-(D) and acceptor (A) fluorophore to characterize conformational states (Figure S1A). Typically, the donor-acceptor pair (DA) is strategically placed within biomolecules to monitor intra-and intermolecular conformational changes and dynamics, by measuring relative changes[14] or absolute distances[19] between both fluorophores. FRET, but also other photophysical strategies, e.g., using (de)quenching or environmentally-sensitive dyes, were shown to be well suitable for the design of protein-based fluorescent sensors[20–25].

Experimental settings that are compatible with FRET encompass most buffer media[26,27] and lysates[28], but even living cells or organelles have targeted more recently[29–33]. This gives access to a wide range of physiological conditions conserving biomolecular structure and function[26,27]. The ability of smFRET to monitor conformational changes and dynamics with sub-nanometer spatial and sub-second temporal resolution has proven to be powerful for studying enzymatic reactions[34–37], nucleic acids[38–40], membrane transport[41,42], molecular motors[43], protein folding[44–47] and many others[3]. In proteins, (covalent) anchor points for the fluorophores are required at specific residues to establish a FRET assay. The most commonly used amino-acid residue for this is a cysteine, which is typically introduced via site-directed mutagenesis and labelling of the resulting thiol with maleimide chemistry[48–51]. Other options such as the introduction of non-natural amino acids[52] or tags[53] exist, yet cysteine maleimide-chemistry remains the gold standard method for fluorophore-labelling on proteins.

Stochastic labelling of proteins, e.g., with two cysteines, is the most common method to introduce the donor and acceptor fluorophore to a protein. This, however, results in a stochastic mix of labeled species since D and A can be linked to either position (Figure S1). While single molecule methods can partially account for the resulting heterogenous mix of labelled biomolecules via advanced experimental schemes (e.g., MFD[54,55], PIE[56] or ALEX[57]; Figure 1B/C, Figure S1B) and data analysis methods[26], the desirable way would be to obtain a homogenous preparation of labelled biomolecules with the desired mix of donor and acceptor dye. Size exclusion chromatography (SEC), which is a common method for the purification of labelled proteins, is generally unable to separate protein with distinct label compositions (Figure S1C), in contrast to ion exchange chromatography (IEX). Here, the selection of buffer, pH or the charge of the fluorophores makes it possible to alter the electrostatic surface potential of a protein and therefore impact its elution behavior[58,59] (Figure S1D). Using this idea site-directed labeling was demonstrated for proteins with two cysteines using a differential labeling approach[60–62,59], via the incorporation of unnatural amino acids[63], or tag-based incorporation of labels[53]. Also, sequential labeling was used to enrich a pure donor-acceptor labelled protein, using distinct reactivities of two protein residues. The latter method was often used in combination with anion exchange chromatography (AIEX) to first separate a single dye species after a first round of labeling with a second separation step via AIEX after a second round of labeling[58].

**Figure 1:**
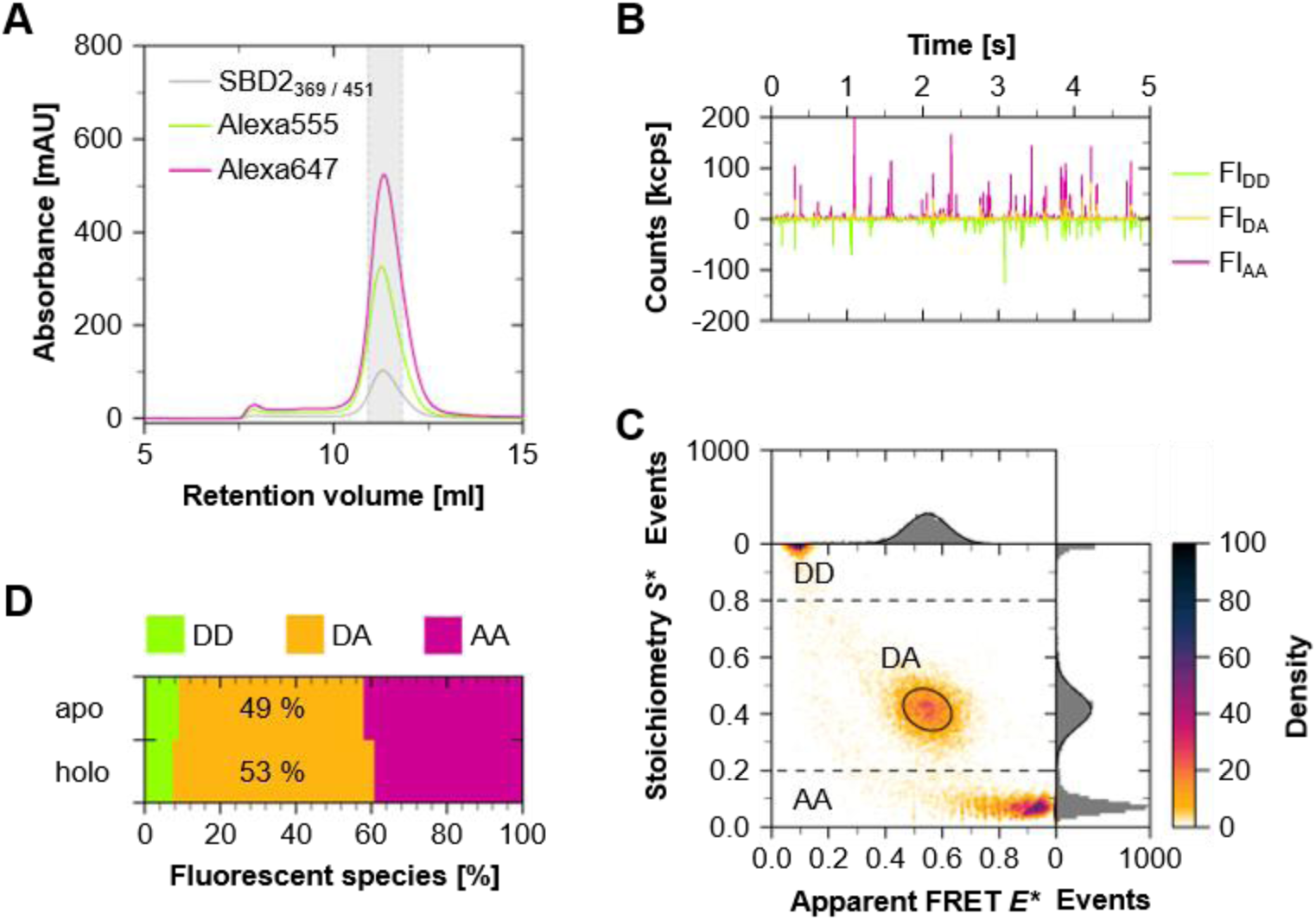
**Characterization of SEC-purified SBD2_369 / 451_ labelled with Alexa555/647**. A: Absorbance profile of the volume dependent SEC elution of SBD2 (280 nm) with Alexa555 (556 nm) and Alexa647 (650 nm) with >90% DOL for both cysteine residues (pooled fraction within the shaded area). B: Fluorescence time traces of the labelled preparation at 100 pM concentration: fluorescence in the donor channel after green excitation (Fl_DD_), acceptor emission after red excitation (Fl_AA_), and sensitized acceptor emission after donor excitation (Fl_DA_). C: *E*S\**-histogram of ligand free (apo-state) resolving donor-only (*S\** > 0.8), acceptor only (*S\** < 0.2), and donor-acceptor species (*S\** = 0.2 - 0.8). D: Relative population of labelled species according to ALEX sorting in both the apo-and holo-(100 µM L-glutamine) conformation.

Here, we present an experimental strategy that combines a stochastic labeling protocol with a single step purification of a protein using AIEX. While AIEX has been used before to purify biomolecules for FRET-assays[58,59], we provide a conceptual basis to systematically design a purification protocol for an arbitrary choice of fluorophores. By first characterizing the interaction of hydrolyzed fluorophore-maleimides with the AIEX-column material, one is able to select pairs of fluorophores that are likely to allow successful purification of donor-acceptor-labelled proteins via AIEX. We demonstrate the strength of the approach with bulk and single-molecule FRET studies of various bacterial substrate-binding proteins. By direct comparison of data from SEC and AIEX we show a significantly improved data quality of proteins purified with the new method.

## RESULTS

We selected three substrate-binding proteins (SBPs), the GlnPQ substrate binding domain 2 (SBD2) from *L. lactis*, maltose binding protein (MalE), and phosphate binding protein (PBP) from *E. coli* for the characterization of the AIEX-based purification approach. These proteins exhibit similar folding patterns featuring two rigid domains connected by a flexible hinge, forming substrate-specific binding sites at their interfaces. Based on crystallographic data of both apo-and holo-conformations^1^, four double-cysteine mutants were selected that showed large changes from long to short inter-dye distances upon ligand binding (SBD2_369 / 451_[64], MalE_36 / 352_[65], MalE_87 / 186_[66,67], and PBP I76G_3 / 86_[68].

We started with a popular pair of dyes, which is used frequently[68] for single-molecule FRET studies of proteins (Alexa Fluor 555 and Alexa Fluor 647). Figure 1 shows a SEC purification of SBD2_369 / 451_ with these dyes. For smFRET analysis we collected fractions from the monodisperse elution peak (Figure 1A), where the absorbances of Alexa Fluor 555 (Alexa555) and Alexa Fluor 647 (Alexa647) coincided with the UV-absorbance of the protein (Figure 1A, shaded area). In these fractions the degree of labelling, DOL, was 0.86 for donor and 0.94 for the acceptor. To characterize the labelling stoichiometry on the protein, we performed microsecond-timed alternating laser excitation (µsALEX[57]) with diffusing molecules. The experiments with dilute protein samples (∼100 pM) directly revealed the heterogeneous nature of the sample, which is seen qualitatively in the fluorescence signals (Figure 1B) and in quantitative burst analysis (Figure 1C/D). A standard all-photon burst search was used for the determination of apparent FRET efficiency (*E\**) and stoichiometry (*S\**) of individual molecules, which could be classified into three distinct populations: D-only (*S\** = 0.8-1.0), DA (*S\** = 0.2-0.8), and A-only (*S\** = 0.0-0.2). When we quantified the number of molecules in each population, we observed that stochastic labelling and SEC purification had a fraction of ∼50% of donor-acceptor labelled protein (Figure 1D). This represents the maximum value for stochastic labelling of two equally reactive binding sites and fluorophores[69,70].

We next tested the ability of a single AIEX purification step to improve the smFRET data quality via separation of protein species with distinct fluorophore stoichiometries (Figure 2). The AIEX conditions were chosen such that interactions between the protein and the column, as well as the fluorophore and the protein were minimal. Continuum electrostatic predictions of the folded protein surface[71–74] suggested that at pH > 7.5 the different SBPs are close to their isoelectric point (Figure S2A). AIEX preparations of the unlabelled mutant SBPs showed low affinities towards the column media at a pH value of 8.5 (Figure S2B), increasing the influence of any charged fluorophore on the ion dependent retention on the AIEX column medium. Based on this we used a purification protocol with pH > 7.5 and salt concentrations of NaCl between 0 and 400 mM increasing by 7.6 mM / ml.

**Figure 2:**
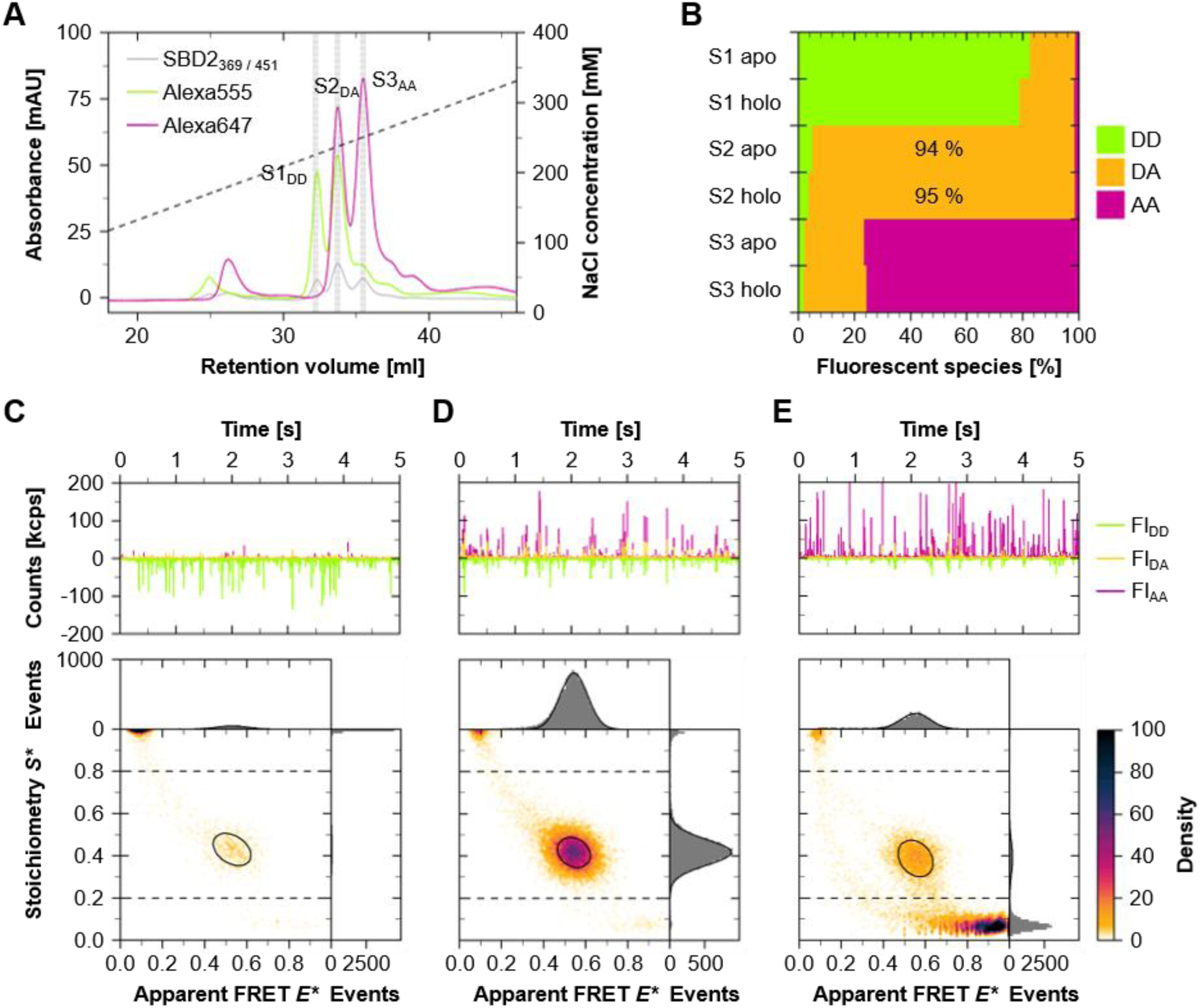
**Characterization of AIEX-purified SBD2_369 / 451_ labelled with Alexa555/647**. A: Absorbance profile of the ion dependent elution of the protein (280 nm) with Alexa555 (556 nm) and Alexa647 (650 nm). B: Relative population of species in the elution fractions S1_DD_, S2_DA_, and S3_AA_ according to ALEX sorting, recorded in the absence (apo) and presence of 100 µM L-glutamine (holo). C – E: Fluorescence time traces at 100 pM concentration and associated *E*S\**-histograms recorded in apo-conformation for species fractions S1-S3.

The resulting chromatogram of SBD2 showed regions of proteins with a single label (Figure 2A, starting at 24 ml) or two fluorophores (Figure 2A, starting at 32 ml). Samples taken from the early elution times, as well as S1_DD_ and S3_DD_ contained predominantly donor-or acceptor-only species, seen in the time traces and corresponding ALEX histograms (Figure 2C, E). Strikingly, the intermediate fraction S2_DA_ had up to 95% of donor-acceptor containing molecules (Figure 2D/B). As a functional control of this fraction, we performed a three-step titration with L-glutamine, i.e., the apo-(without ligand), holo-(with 100 µM) and K_D_-value (1 µM[64,65,75]), confirming the functionality of the sample after AIEX purification (Figure S3).

To assess the positive impact of the AIEX-purification on the statistics and measurement times of FRET assays, we prepared three SEC purified samples of SBD2 with Alexa555/647 that had average DOL values of 34%, 61%, and 90% for each of the two cysteine positions (Figure S4A). Using ALEX, we found that a DOL ∼34%, DOL ∼61%, and DOL ∼90% relates to 26%, 41%, and 49% of donor-acceptor-labelled protein, respectively (Figure S4B). With these reference numbers we investigated what time and burst numbers are required to reliably fit data of SBD2 via 2D Gaussians for data sets in which both apo-and holo-conformations were equally populated (Figure 3A-C).

**Figure 3:**
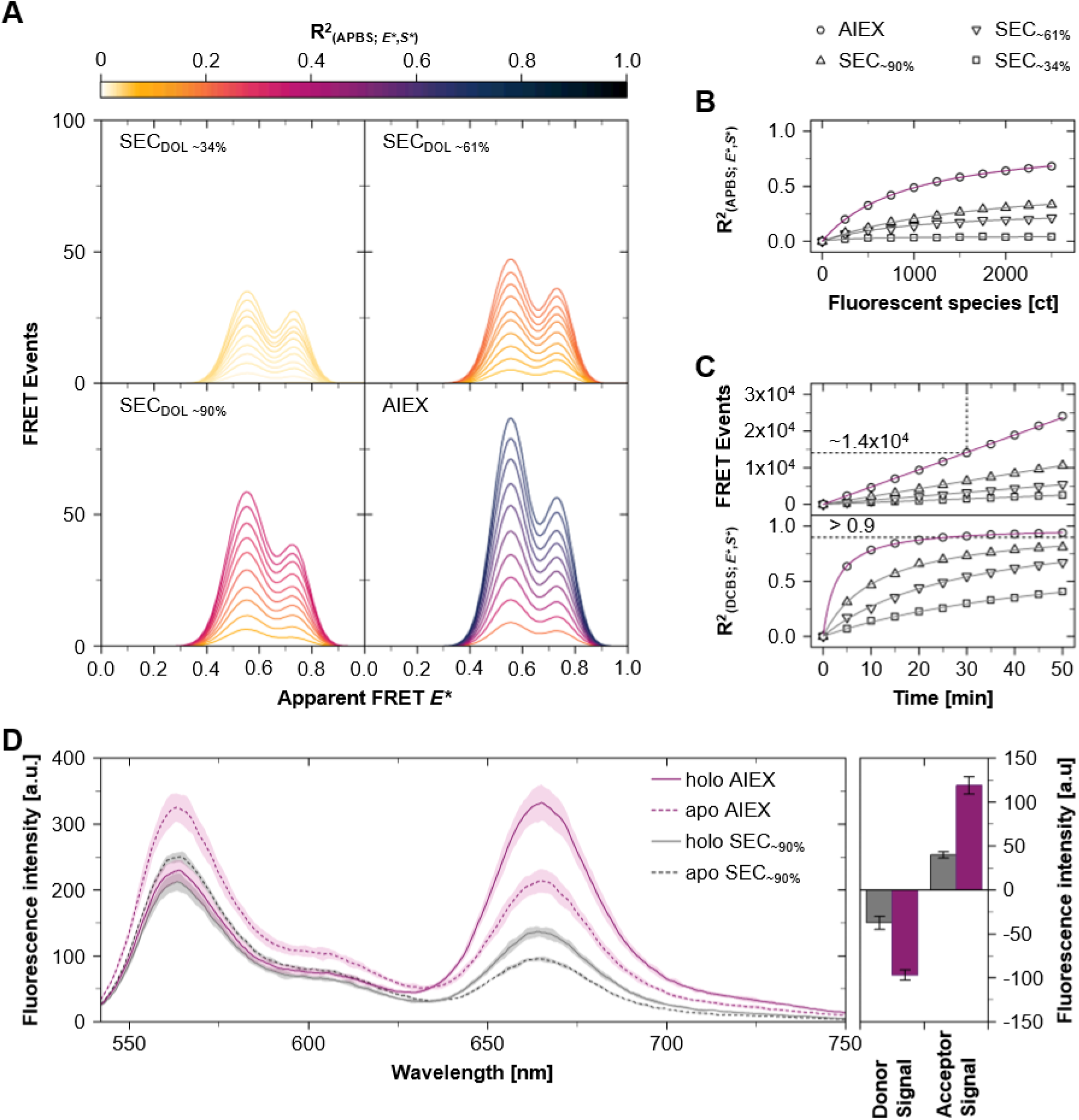
Quantitative analysis and comparison of SEC and AIEX purified samples. A: Fit curves of FRET data of labelled SBD2_369 / 451_ at 1 µM L-glutamine recorded in increments of 250 detected molecules using fixed apo-and holo-states. B: R^2^as a function of the total (APBS) fluorescent species count. C: Increase in the number of the total FRET events and improvement of R^2^ for a 2D (*E\**-*S\**) Gaussian fit as a function of time. D: Mean fluorescence emission intensity of SEC and AIEX purified SBD2 with Alexa555/647 using excitation at 532 nm (n = 3; technical repeats with stanard deviation). The apo-to holo-state transition is indicated by the decrease in donor-(563 nm) and the increase in acceptor emission (665 nm) after addition of 100 µM L-glutamine.

For this assessment, we ran an all-photon burst search and plotted the fits to an *E\**-histogram for all events in the *S\** region between 0.2-0.8 for every 250 detected molecules (Figure 3A). It was apparent that increasing DOL from SEC-purified protein yielded a higher number of bursts with FRET information (Figure 3A). Also, the goodness of fit described by R^2^ (Figure 3A/B) improved much faster, particularly for AIEX samples in relation to the total number of detected molecules (Figure 3C). Using AIEX we were able to obtain >10.000 burst events with donor-acceptor species in less than 30 minutes which gave values of R^2^ > 0.9 (Figure 3C).

Furthermore, we performed comparative bulk FRET measurements of AIEX and a SEC purified sample with a DOL of ∼90%. The fluorescence spectra after donor excitation showed characteristic emission peaks for Alexa555 and FRET-sensitized emission of Alexa647 (Figure 3D). Upon addition of saturating ligand concentrations an increase of FRET efficiency was observed by the anti-correlated changes of donor-and acceptor-emission with a drastic improvement of the signal amplitude for the AIEX sample (Figure 3D). We note that the ion exchange column type (MonoQ 5/50 GL or Capto HiRes Q 5/50) did not affect the quality of the preparation (Figure S5).

Motivated by the significant improvement of data quality in both single-molecule and bulk assays (Figure 3), we reasoned that a characterization of different dyes would be helpful to establish a systematic purification approach using AIEX. For this we considered rhodamine-and cyanine-dyes with varying number of charges and charge distributions (Figure 4, Figure S6A). In buffer solutions at pH values between 7.5 – 8.5 all dyes are supposed to carry one positive charge and additional two (Alexa532, sCy3, sCy5), three (Alexa488, Alexa568), or four negative charges (AF555, AF647, Alexa555, Alexa647) due to deprotonation of the attached sulfonate and carboxyl groups (Figure 4, Figure S6A).

**Figure 4:**
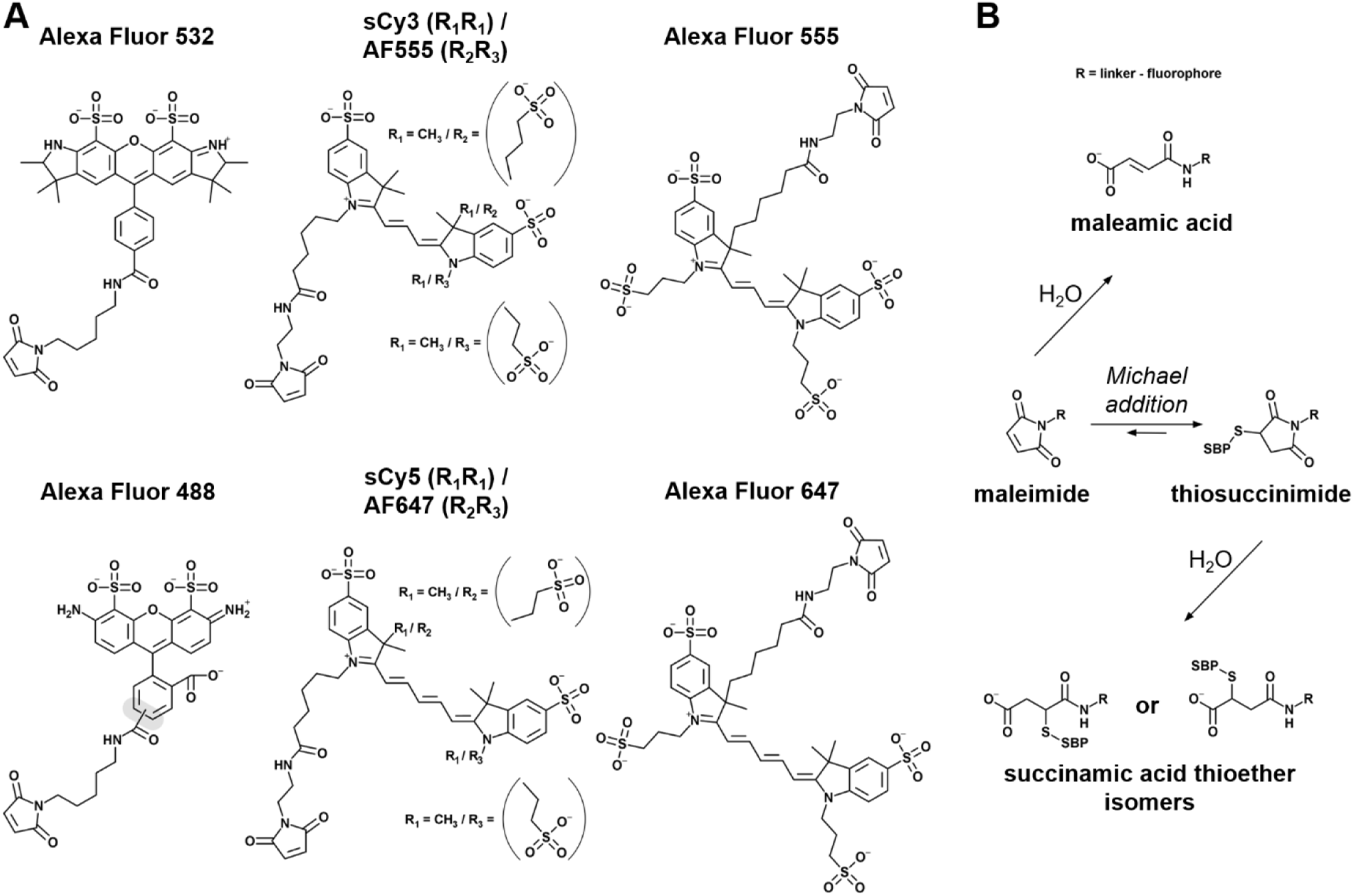
Chemical information on fluorophores and labelling chemistry. A: Chemical structures of anionic fluorophore maleimides: Alexa532 (Thermo Fisher Scientific[80]) and Alexa488 (Thermo Fisher Scientific[81]), as well as cyanine fluorophores sulfo-Cyanine3 (sCy3; Lumiprobe[82]), sulfo-Cyanine5 (sCy5; Lumiprobe[83]), AF555 (Jena Bioscience, literature[66]), AF647 (Jena Bioscience, literature[66]), Alexa555 (Thermo Fisher Scientific, literature[66]), and Alexa647 (Thermo Fisher Scientific, literature[66,84]). At the time of submission, no structural information was available for ATTO643 maleimide and only a net charge number of-1 after coupling was provided by the manufacturer. Additionally, ATTO643 is supplied as a mixture consisting of three isomers with negligible differences regarding their spectral properties. Similarly, Alexa488 and Alexa568[85] are provided as two isomers where the linker attachment at the central phenyl-ring can occur at both the meta-or para position in relation to the fluorophore core. B: Fluorophore-SBP bioconjugation via Michael addition and the hydrolytic reaction scheme of the thiosuccinimide intermediate and maleimide group according to ref.[50].

Bioconjugation of the fluorophore maleimide to a protein-thiol via a Michael Addition and subsequent irreversible hydrolysis of the thiosuccinimide intermediate results in the formation of stable succinamic acid thioethers[76–79,49,50] (Figure 4B). We thus reasoned that the state of the protein-coupled succinamic acid thioether fluorophore is similar to a hydrolyzed fluorophore maleamic acid (Figure 4B). Consequently, interactions of the hydrolyzed dye with the column material should allow predictions for the feasibility of fluorophore-dependent AIEX purification of labelled protein species. As expected[58] the gradual elution (7.6 mM NaCl/ ml) of the (partially) hydrolyzed fluorophores yielded a fluorophore specific elution pattern with a distinct separation for mixtures of different dye molecules (Figure 5, Figure S6-8). The hydrolyzed form of the dye showed an overall higher affinity than the respective maleimides (Figure S6B, S7A/C, S8).

**Figure 5:**
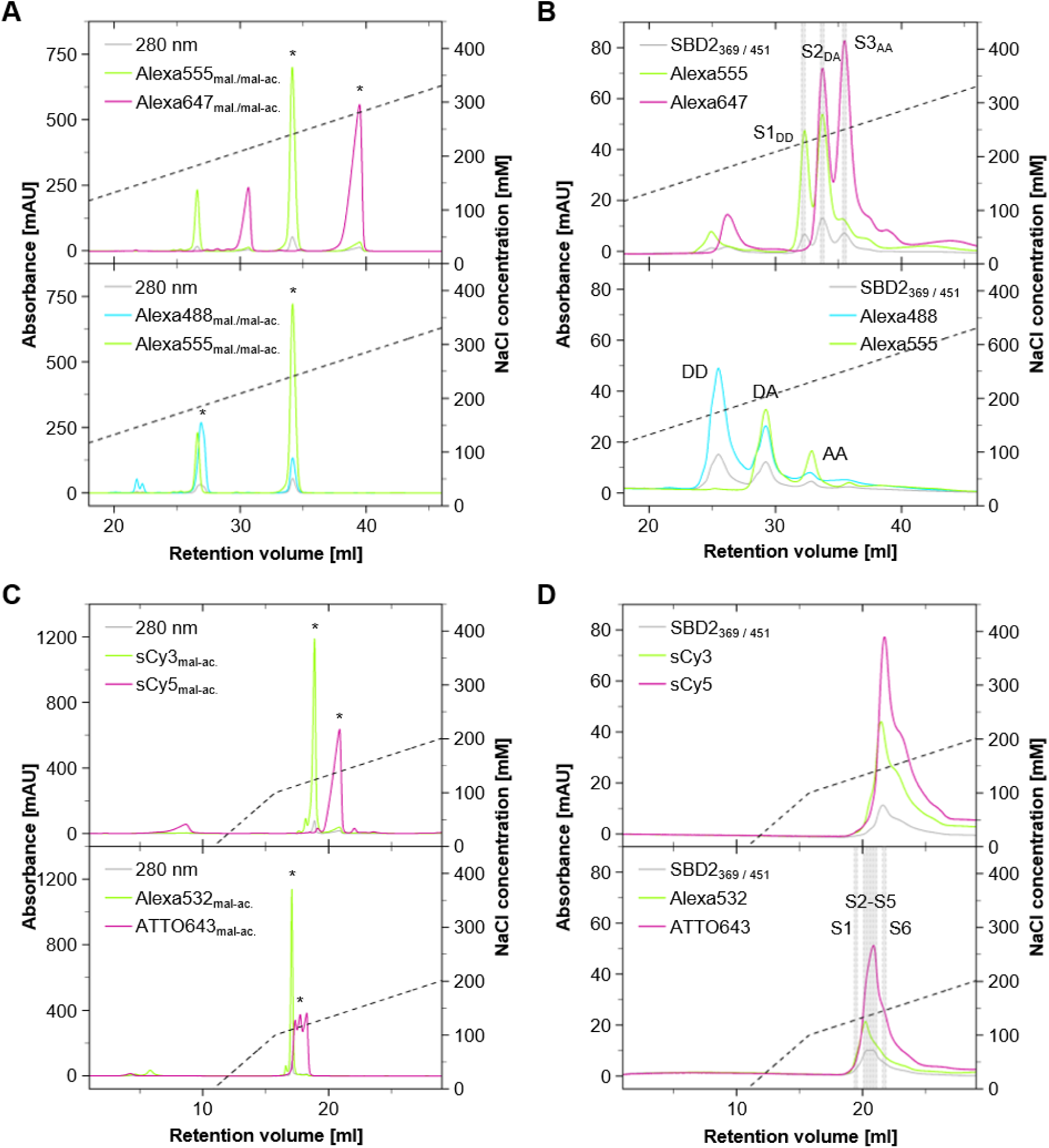
Side-by-side comparison of fluorophore elution and labelled proteins on AIEX. A/C: Absorbance profiles of the ion dependent AIEX elution (pH 7.5, 7.6 mM NaCl/ ml) of the partially hydrolyzed (maleimide/mal. and *maleamic acid/mal-ac.) fluorophore pairs Alexa555/647 (556/650 nm), Alexa488/555 (493/556 nm), sCy3/sCy5 (548/646 nm), and Alexa532/ATTO643 (528/643 nm) using 14 nmol per fluorophore. B/D: Ion concentration dependent elution profiles (pH 8.5, 7.6 mM NaCl/ ml.) for the SBD2 protein (280 nm) labeled with respective fluorophore pairs.

As expected for the well-separated AIEX-elution profile of SBD2 with Alexa555/647 (Figure 2A), hydrolysis of these two fluorophores and AIEX analysis showed large differences in their retention volume of the isolated dyes (Figure 5A). Combinations of Alexa488 with either Alexa555 (Figure 5A) or Alexa647 (Figure S7A) showed similar differences. Importantly, these differences had predictive character for the resulting separation of labelled protein species with the two dye pairs (Figure 5B, Figure S7B). In contrast, a combination of sCy3/sCy5 or Alexa532/ATTO643 showed a distinct pattern: the maleimide fluorophores already eluted prior to the onset of the ion gradient and association of isolated dyes to the column occurred only for the hydrolyzed fraction of the dyes (Figure 5C, Figure S8B).

But the differences in elution volumes for both combinations were minor. Consistent with these observations the protein preparations did not allow to separate the different labelled species via AIEX (Figure 5D). In both cases AIEX of SBD2 with sCy3/sCy5 and Alexa532/ATTO643 did not effectively separate labelled species (Figure 5D) and only showed a maximum of ∼60% donor-acceptor containing protein species (Figure S9).

We finally tested our strategy with double-cysteine variants of the maltose-and phosphate binding protein (Figure 6). We chose two distinct variants of MalE with residues with low (MalE_87 / 186_) and high tendencies (MalE_36 / 352_) for fluorophore interactions based on reported fluorescence anisotropy values[65–67]. According to expectations MalE_87 / 186_ followed the same pattern seen in SBD2 when labelled with Alexa555/647. As shown in Figure 6A we were able to separate the DA fraction from the other doubly-labelled species. Notably, higher starting concentrations of salt (∼100 mM NaCl) were beneficiary to enrich the DA-labelled species to >97% (Figure 6A/B, Figure S10) due to different surfaces charges of MalE compared to SBD2 (Figure S11). MalE_36 / 352_ showed a distinct pattern with five different peaks (S1-S5) containing doubly-labelled proteins at elution volumes >23 ml (with 100 mM NaCl starting concentration, Figure 6C) or >35 ml (without NaCl addition, Figure S10). Considering the DOL and ALEX experiments (Figure 6D) we can assign S1 to donor-donor, S4&S5 to acceptor-acceptor labeled species and S2&S3 to proteins with both donor and acceptor. Since strong fluorophore-protein interactions have already been reported for position 352C in MalE[66,67], it is likely that the separation of S2 and S3 originate from proteins with donor and acceptor at either cysteine position: Alexa555-36C / Alexa647-352C, Alexa647-36C / Alexa555-352C.

**Figure 6:**
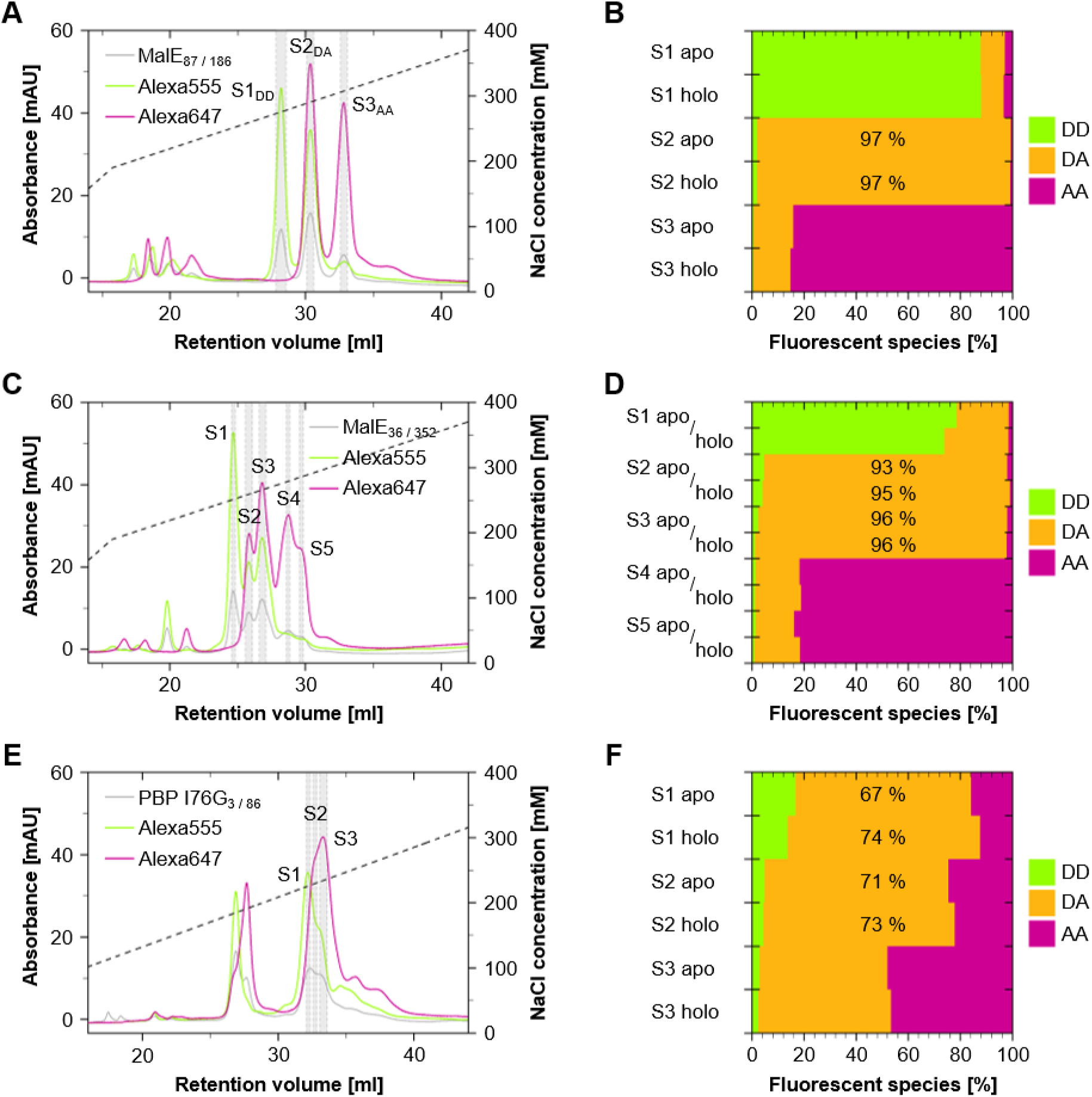
AIEX purification and ALEX characterization of different protein variants with Alexa555/647. A, C: Absorbance profiles of the ion-dependent separation (pH 8.5, 7.6 mM NaCl/ ml, initial supplementation with 100 mM NaCl) of the different protein variants: MalE_87 / 186_ and MalE_36 / 352_ protein (280 nm) with Alexa555 (556 nm)/647 (650 nm). B, D: Relative populations of labelled protein species based on ALEX. E/F: Similar data for PBP.

AIEX of PBP I76G_3 / 86_ Alexa555/647 did not result in a similar separation of the double labelled species even though the single labelled fractions eluted at lower ion concentrations and appeared as well-defined peaks around 26-27 ml (Figure 6E). Variations in pH-value and the initial supplementation of the AIEX buffer with 100 mM NaCl were tested, but did not improve the overall pattern of the AIEX elution profile of PBP (Figure S12). In between S1-S3 the ratio of donor to acceptor fluorophore (D/ A ratio) gradually shifted from favoring a donor-to acceptor excess (Figure 6E/F). Despite the cutback, both S1 and S2 contained >70% donor-acceptor-labelled protein for PBP, and we performed a functional check of the protein with these fractions via a ligand titration. As for all SBPs in Figure 6, we found a ligand dependent transition from the unliganded apo-to holo-state seen as a mix of apo-and holo-state at concentrations close to the K_d_ values of each protein (SBD2_369/451_: 1 µM L-glutamine[64,65,75], MalE_36 / 352_/ MalE_87 / 186_: 1.5 µM D-maltose[86,87,65–67], PBP I76G_3 / 86_: 10 µM phosphate[68]). The measurements confirmed that the AIEX procedure did not impair the functional integrity of the SBPs (Figure S3). In PBP we always found a much higher protein content with a single label relative to fractions with two (Figure 6) in comparison to SBD2 and MalE. Potentially, this ratio of singly-labeled to doubly-labeled species might help to identify whether the labeling of a second cysteine after initial labeling of the first position is hindered by the presence of a primary attached fluorophore molecule.

## DISCUSSION AND CONCLUSION

Based on AIEX we here present a strategy for stochastic labeling of protein residues with donor and acceptor dyes for FRET experiments. For rationale design of labelling protocols beyond our own work, we provide a conceptual basis by characterizing the interaction of fluorophore-maleimides with the AIEX-column material. Based on this approach we are able to select (and predict) which pairs of fluorophores allow successful purification of donor-acceptor-labelled proteins via AIEX using various bacterial substrate-binding proteins as representative examples. We also found our approach to work with different AIEX-columns, since we observed no major differences between the Capto HiRes Q 5/50 and MonoQ 5/50 GL (the latter is discontinued; comparison see Figure S5, S13).

We believe that not only the presented results, but also previous usage of similar purification workflows support the usefulness of our single step purification protocol. We obtained high-quality protein preparations via an AIEX purification in which conventional SEC purification did not yield a sufficiently high fraction of donor and acceptor labelled proteins. Examples include MalE_31 / 212_ used for absolute intramolecular distance determination via smFRET and aSAXS[88] and for optimal labelling of SBD2 for smFRET studies using blue-green dyes[89]. Significant improvements are also expected for bulk applications where the removal of inactive sensor fractions would substantially increase the dynamic range of assays as shown in Figure 3D. This is not only true for FRET, but would also apply to a broad range of biosensors that utilize different photophysical strategies to monitor conformational changes such as (de)quenching[23] of two identical fluorophores or environmentally-sensitive dyes[24].

Based on our results, we can summarize the considerations that are relevant to design a successful purification protocol for unknown protein systems. We found that protein stability and an overall minimized surface charge aid in the AIEX purification. Yet, smaller variations in pH did not significantly improve the AIEX profile of Alexa555/647 on SBD2 (Figure S14B) or PBP (Figure S12), which can be seen in the pH independent elution of hydrolyzed Alexa555/647 (Figure S8C). These findings suggest that the fluorophore (charge) is probably more relevant than pH dependent charges of the protein (Figure S2B). Thus, it may not be necessary to work at pH values close to the isoelectronic point of the respective protein for AIEX purification (Figure S2A).

To explain the distinct elution profiles of SBD2, MalE and PBP we considered both the location and density of surface charges on the protein (Figure S11). Continuum electrostatic predictions[71–74] of the folded protein surfaces suggest net negative charges at pH 8.5 for SBD2, MalE, and PBP (Figure S2A). Yet, AIEX purification of PBP did not yield a clear separation of the doubly-labelled species. We also observed clear differences in the AIEX-profiles of MalE_36 / 352_ and MalE_87 / 186_. The latter may arise from the fact that fluorophores can homogenously screen positive charges in MalE_87 / 186_ at either position, while surface charges show a clear position dependency in MalE_36 / 352_ (Figure S11D). In case of PBP I76G_3 / 86_ a number of positive charges were found to be in and around the binding pocket for phosphate, however, volumetric calculations[90] suggested that the accessible volumes (AVs) of neither Alexa555 nor Alexa647 were overlapping with the positive surface areas (Figure S11B), which might interact strongly with the anionic fluorophores. The only difference we were able to spot was that PBP has a single patch of positive residues in the AVs (Figure S11B) absent in SBD2 (Figure S11A), while MalE has an overall positive surface charge (Figure S11C/D). This provides, however, no clear line of argumentation to explain the observed effects.

We also considered whether the N-terminal His_10_-Tag, which is located within the AV of the SBD2 451C position, can impact the interaction of the protein with the AIEX matrix, e.g., due to charge screening. However, for SBD2_369 / 451_ in combination with Alexa555/647 we observed that the overall elution pattern was maintained even after the enzymatic removal of the His_10_-Tag (Figure S14A). In summary, a qualitative inspection of surface charges to explain differences in AIEX-profiles seems at best to provide a handwaving line of reasoning for differences in the AIEX profiles.

What about the chemical properties of the fluorophores? Overall, the number of charges of the dyes served as a good predictor for the retention on the AIEX column medium. We also observed that increasing the distance between negatively charged sulfonate groups generally enhanced the affinity towards the AIEX medium (see Figure 4 and 5). The comparison of constitutional isomers of the cyanine dyes Alexa555 / AF555 and Alexa647 / AF647 showed that the isolated hydrolyzed dyes had nearly identical AIEX-profiles, while for covalent attachment to SBD2_369 / 451_ we obtained only clearly separable DA species for Alexa555/647 but not for AF555/647. To us this suggests that the concrete charge distribution of the dye on the protein has high relevance for dye-protein interactions (also observed in anisotropy decays of AF555 vs. Alexa555 on residue 352 in MalE[66]) leading to the observed differences in AIEX behavior for the unbound and bioconjugated fluorophore (Figure S7C/D). In summary not only the number of charges of the fluorophore determines its affinity for the AIEX column material, but also the steric distribution of the charged sulfonate and carboxyl groups in particular in its bioconjugated form.

Considering the impact that even such small variations in the chemical structure of the different fluorophores have on AIEX elution profiles, ion exchange methodology may aid in the development of new fluorophore conjugates using alternative linkers or additional functional groups[91]. Additionally, AIEX could help to remove certain fluorophore sub-populations (e.g. isomers) prior to labeling, that could otherwise lead to unwanted labelled protein species[59]. Furthermore, we believe that our work highlights the relevance of obtaining fluorophore structures and the importance of a uniform nomenclature for fluorescent molecules used for bioconjugation, including steric information about functional groups not only in the realm of integrative structural modelling using FRET.

A final consideration is the question is how dye protein binding impacts photophysical behavior. AIEX cannot mitigate issues which are normally tackled by synthetic modifications of the fluorophore core[92] or the (covalent) addition of photoprotective agents and oxygen scavenging systems[93–101]. Yet, AIEX can increase the photophysical homogeneity, help to identify whether dyes interact with the protein surface and thus help to understand which amino-acid residues get into contact with it (see comparison of the MalE variants in Figure 6). Additionally, the effects of thiol conjugation on photophysical properties have been described[101] and our data suggests that AIEX can discriminate between minute structural differences in the dyes (Figure 2 vs. Figure S7) and labelling positions (Figure 6A vs. 6C). Possible future improvements could be the prevention of the ring opening after Michael-addition via hydrolysis[49,50], which could be a mechanistic factor that promotes the photobleaching (“thioether effect”) that was described for multiple cyanine and rhodamine fluorophores[101].

## METHODS

All methods used in this paper are provided in Supplementary Note 1 since most of them were described in detail previously[64,65,102]. We provide details on the AIEX-based protein labelling procedure here:

### Fluorophore labelling of SBD2, MalE, and PBP

The stochastic maleimide labelling and purification followed an already established protocol[64,65,102]. For each labeling reaction 600 µg protein from stocks stored at-80°C were used. SBD2, MalE, and PBP variants were incubated for one hour in the respective labelling buffer (SBD2: 50 mM Tris-HCl pH 7.6, 150 mM NaCl/ MALE: 50 mM Tris-HCl pH 7.4, 50 mM KCl/ PBP: 20 mM Tris-HCl pH 7.5, 100 mM NaCl)) supplemented with 1 mM DTT to retain the reduced state of the cysteine residues. Proteins were then immobilized by metal affinity on a Nickel functionalized agarose medium (Ni^2+^-Sepharose 6 Fast Flow, Cytiva). Then the maleimide labelling reaction was carried out in the protein specific labelling buffer overnight at 4°C with 25 nmol of each fluorophore. Subsequently, the resin-bound proteins were washed with the respective labelling buffer and eluted with 500 µl labelling elution buffer (MalE: 50 mM Tris-HCl pH 8.0, 50 mM KCl, 500 mM imidazole/ SBD2: 50 mM Tris-HCl pH 7.6, 150 mM NaCl, 500 mM imidazole/ PBP: 20 mM Tris-HCl pH 8.0, 100 mM NaCl, 500 mM imidazole).

### Labelled protein purification by liquid chromatography (AIEX or SEC)

The labelled MalE, SBD2 and PBP were prepared for further purification by removal of remaining salts and imidazole from the labelling elution buffer that could otherwise interfere with the anion exchange process using a Sephadex G-25 medium (PD MiniTrap G-25, Cytiva). The labelled SBPs were eluted stepwise in a total of 1 ml anion exchange sample buffer (10 mM Tris-HCl pH 7.5-8.5). The anion exchange column (ÄKTA pure chromatography system, Cytiva; MonoQ 5/50 GL column, Cytiva or Capto HiRes Q 5/50, Cytiva) was set up with a 5-column volume H_2_O_dd_ wash, a 10-column volume equilibration with anion exchange sample buffer, 10-column volume equilibration with anion exchange elution buffer (10 mM Tris-HCl pH 7.5-8.5, 1 M NaCl) and a final 20-column volume equilibration with anion exchange sample buffer. Labelled SBPs (500 µl pooled fractions with the highest concentration) were loaded onto the column, and the resin-bound protein was washed with 10 column volumes anion exchange sample buffer. For the consecutive elution a linear increase of elution buffer to sample buffer ratio with a slope corresponding to 7.6 mM NaCl/ ml was chosen (MonoQ 5/50 GL: 5.1 mM NaCl/ min for PBP). All steps were done with a 0.5 ml/ min flow rate. Different fractions containing fluorescently labelled species of SBD2, MalE, and PBP variants were selected according to their absorbance values. Labeling efficiencies and protein concentrations of the isolates were estimated based on the protein molar extinction coefficients and the expected parameters of the conjugated fluorophores (Alexa488: ʎ_exc_ = 493 nm; 72.000 l*mol^-1^*cm^-1^/ Alexa555: ʎ_exc_ = 556 nm; 158.000 l*mol^-1^*cm^-1^/ Alexa647: ʎ_exc_ = 651 nm; 265.000 l*mol^-1^*cm^-1^/ sCy3: ʎ_exc_ = 548 nm; 162.000 l*mol^-1^*cm^-1^/ sCy5: ʎ_exc_ = 646 nm; 271.000 l*mol^-1^*cm^-1^/ Alexa532: ʎ_ex_ = 528 nm; 100.000 l*mol^-1^*cm^-1^/ ATTO643: ʎ_exc_ = 643 nm; 150.000 l*mol^-1^*cm^-1^/ AF555: ʎ_exc_ = 556 nm; 158.000 l*mol^-1^*cm^-1^/ AF647: ʎ_exc_ = 651 nm, ʎ_em_ = 671 nm; 270.000 l*mol^-1^*cm^-1^). Values were corrected considering the respective contribution to the absorbance at 280 nm and to the second fluorophore (if applicable). A purification of SBD2_369/ 451_ labeled with Alexa 555/Alexa647 using size-exclusion chromatography (ÄKTA pure chromatography system, Cytiva; Superdex 75 Increase 10/300 GL, Cytiva), served as a control sample for comparative spectroscopic evaluations. If not stated otherwise, all AIEX procedures in this study were performed using a Capto HiRes Q 5/50 (Cytiva) anion exchange column. Additional data on the MonoQ 5/50 GL (Cytiva, discontinued) as an alternative anion exchange column is provided in Figure S5, S13. The AIEX separation of fluorophore maleimides and fluorophore maleamic acid mixtures were carried out at pH 7.5 in 10 mM Tris-HCl to minimize uncontrolled hydrolysation after the sample preparation (hydrolyzation: 3 h, 25 °C, pH 9.5, 10 mM Tris-HCl).

## ACKNOWLEGDEMENTS

This work was financed by Deutsche Forschungsgemeinschaft (DFG-879CO-4/1 to T.C.), Bundesministerium für Bildung und Forschung (KMU grant „quantumFRET“ to T.C.) and the European Comission (ERC-PoC 101069307 – BIO-LINKER to T.C.). We thank Douglas A. Griffith for the suggestion to use IEX chromatography for the purification of fluorophore-labelled proteins and biosensors.

## AUTHOR CONTRIBUTIONS

N.D.W. and T.C. designed the study and the experiments. N.D.W. performed research, analysed data and prepared figures. T.C. acquired funding and supervised the project. N.D.W. and T.C. interpreted the results and wrote the paper.

## Supplementary Information

## SUPPLEMENTARY DATA

**Figure S1:**
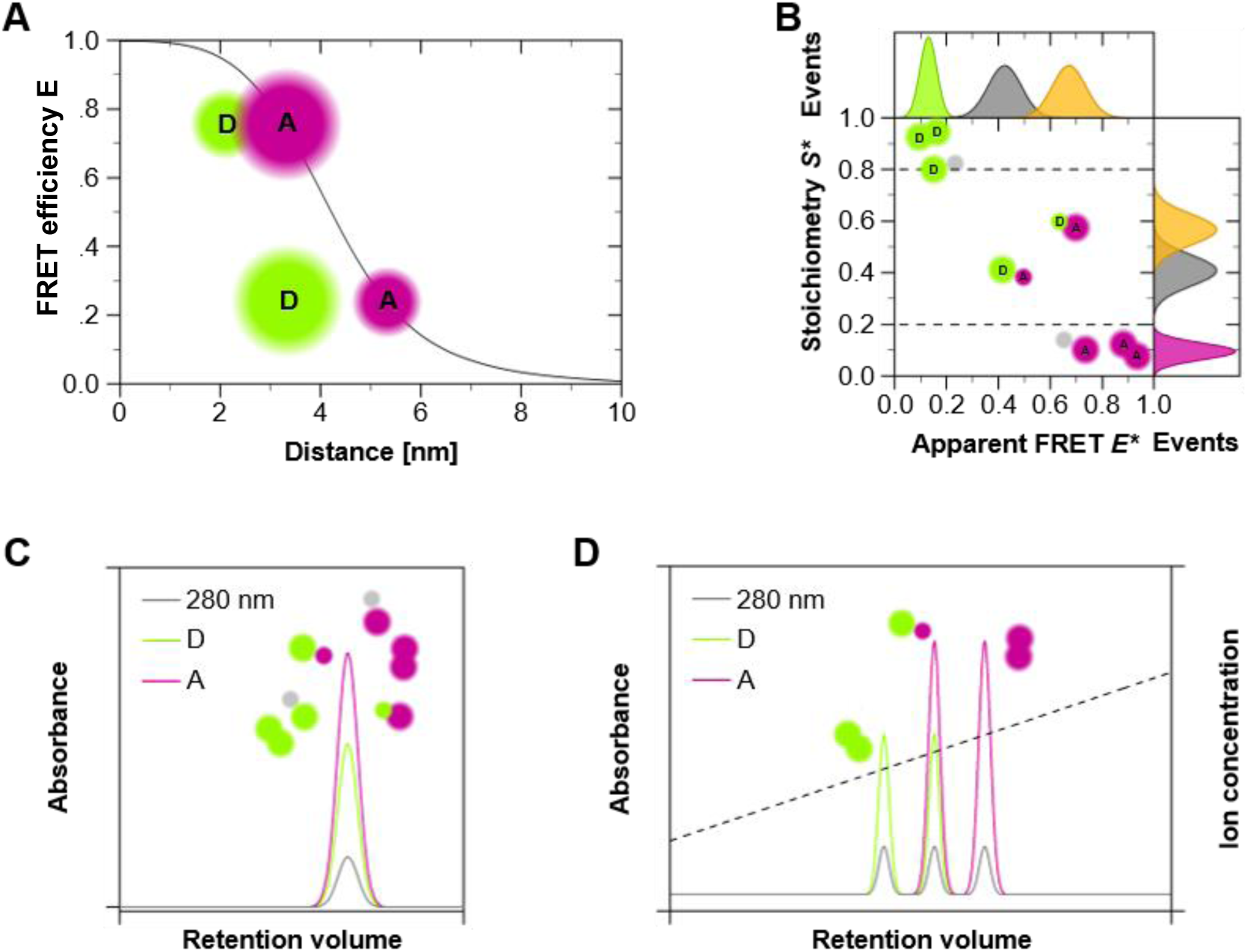
Background on FRET and purification of proteins via FPLC. A: Dependency of accurate FRET efficiency *E* on the absolute distance between a donor (D) and an acceptor (A) dye. B: Setup dependent *E*-S\** Histogram linking Apparent FRET Efficiency *E\** and Stoichiometry *S\** of smFRET measurements using µsALEX. Apparent FRET Efficiency *E\** is defined as the ration between acceptor fluorophore (A) emission and the total emission of acceptor and donor fluorophore (D) after donor excitation. Stoichiometry is defined as D and A emission after D excitation normalized to the emission of both both dyes after D and A excitation allowing to separate species based on brightness: D-only (*S\** > 0.8), A-only (*S\** < 0.2), and D and A (*S\** = 0.2 to 0.8). C: Size-Exclusion Chromatography (SEC) profile showing the co-elution of protein (280 nm) and the covalently attached fluorescent labels. D: Ion Exchange (IEX) profile of stochastically labeled (e.g. D and A) SBPs. Diverse fluorophore modifications may alter the affinity of a protein bioconjugate towards the IEX-medium changing the threshold of an ion-dependent displacement during IEX purification. This allows for the species-selective elution of D-only, A-only, and D and A labelled SBPs.

**Figure S2:**
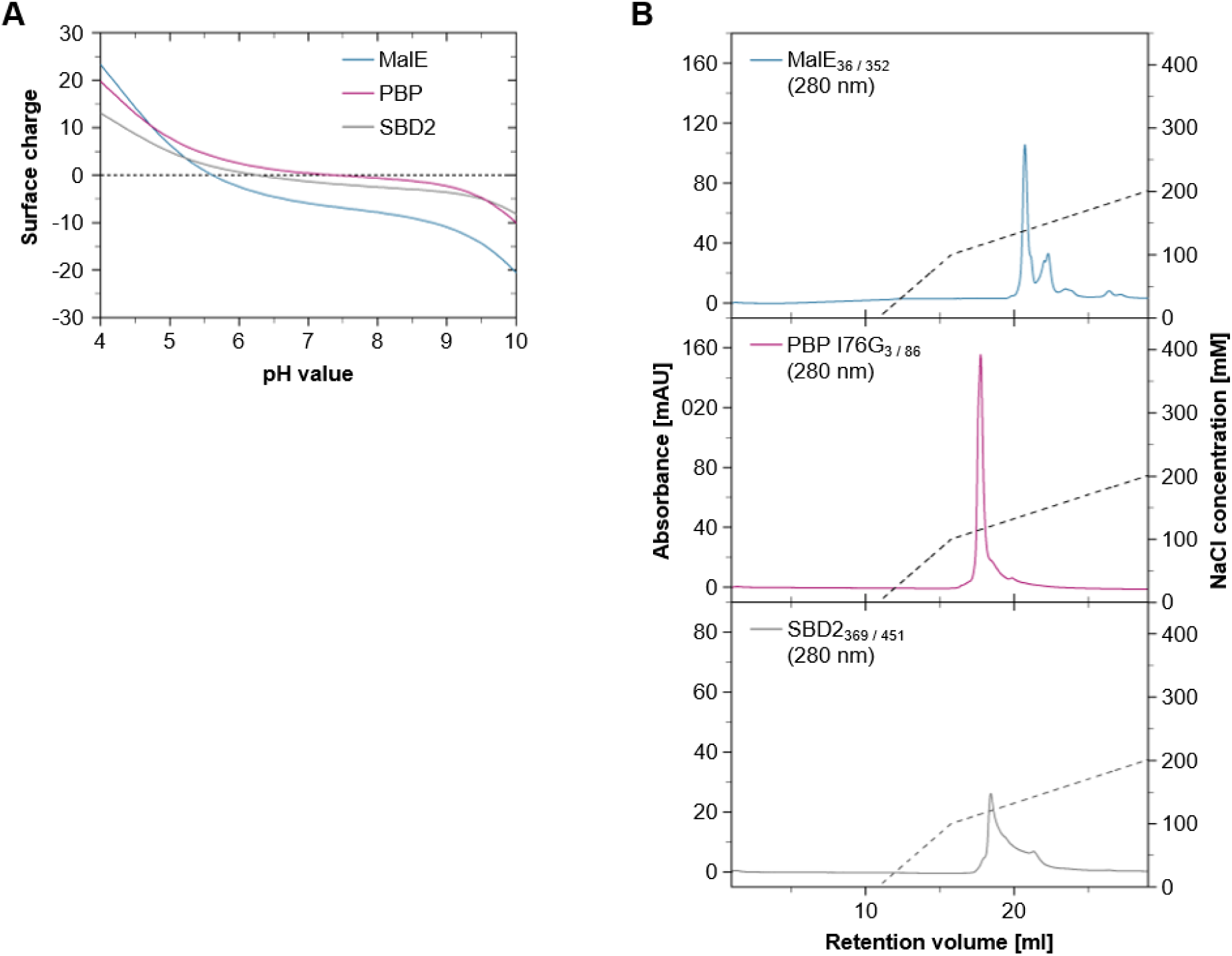
Characterization of the label-free SBP proteins. MalE_36 / 352_, PBP I76G_3 / 86_, and SBD2_369 / 451_, based on their surface charge. A: Continuum electrostatic predictions of the individual folded protein surfaces of MalE, PBP I76G, and SBD2 close to the isoelectric point (IEP). B: AIEX absorbance profiles (Capto HiRes Q 5/50) of the ion dependent purification (pH 8.5, 7.6 mM NaCl/ ml) of the unlabeled SBPs. Absorbance determined at 280 nm.

**Figure S3:**
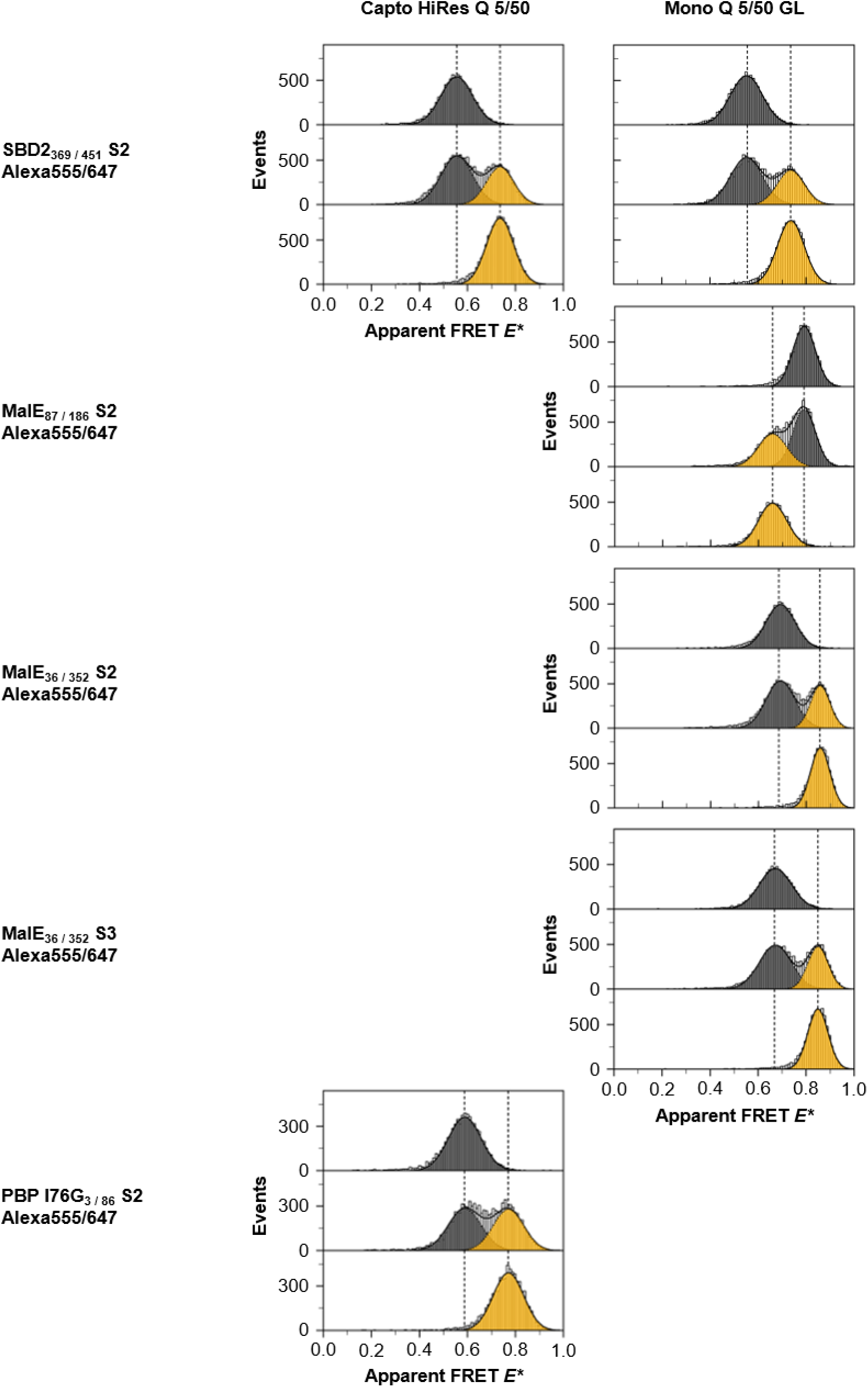
Mini titrations of substrate binding protein. 2D Gaussian fitted (*E*\*-*S*\*) FRET populations visualize the apo-(grey) to holo-(yellow) state transition, with fits using fixed values of mean E* and S* known from apo and holo conditions for Capto HiRes Q 5/50 or MonoQ 5/50 GL. The data used distinct concentrations of ligand for SBD2_369 / 451_, (100 µM / 1 µM L-glutamine), MalE_36 / 352_ (1 mM / 1.5 µM D-maltose), MalE_87 / 186_ (1 mM / 1.5 µM D-maltose), and PBP I76G_3 / 86_ (500 µM / 10 µM phosphate).

**Figure S4:**
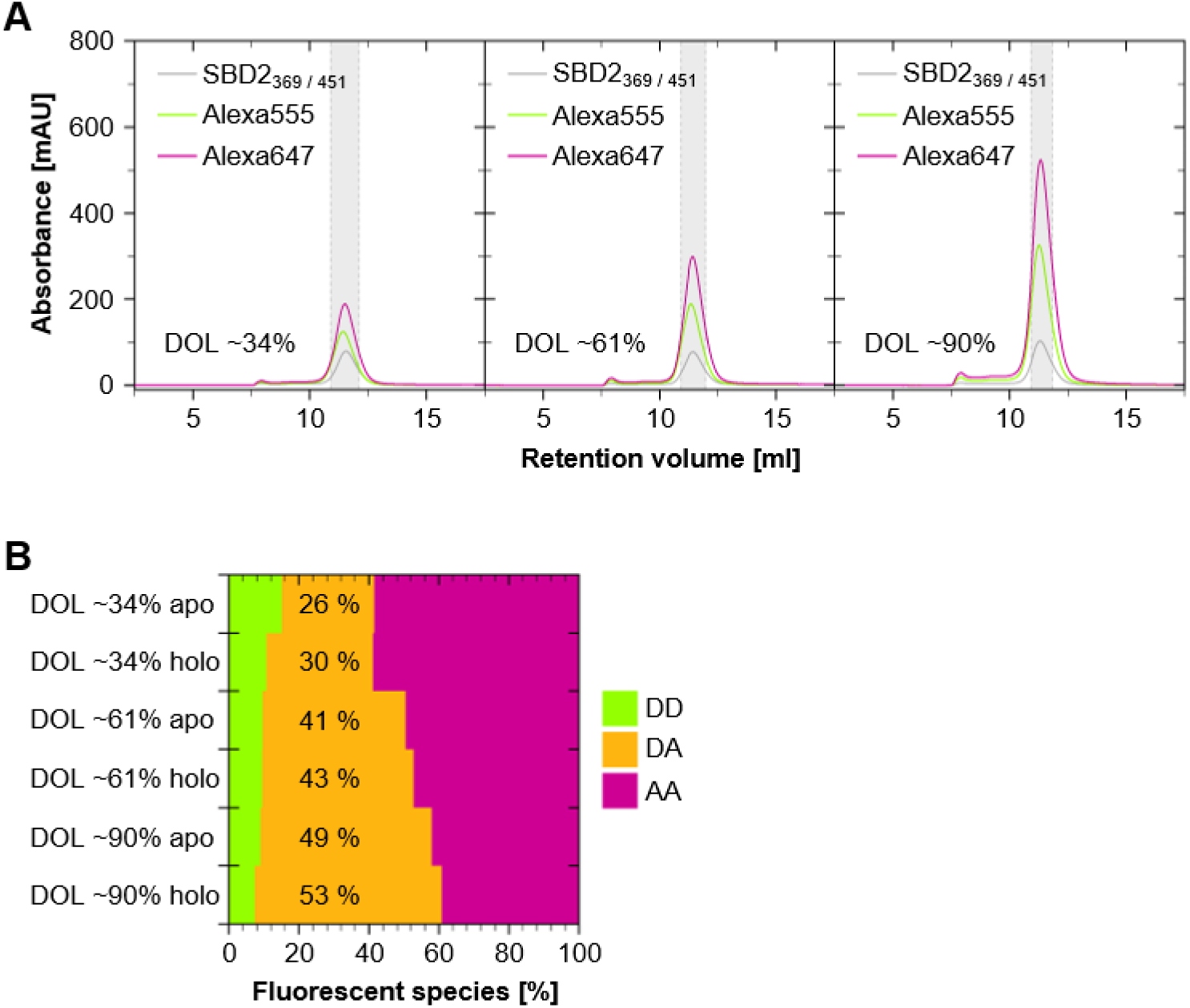
Characterization of SEC purified SBD2_369 / 451_ with Alexa555/647. A: Three individual absorbance profiles of the volume dependent SEC elution of SBD2_369 / 451_ (280 nm) labelled with Alexa555 (556 nm) and Alexa 647 (650 nm) with average ∼34%, ∼61%, and ∼90% DOL for both maleimide receptive cysteine residues. Estimations are based on the main labelled protein fraction (shaded area). B: Relative population of DD, DA, and AA labelled species within the SEC_DOL ∼34%_, SEC_DOL ∼61%_, and SEC_DOL ∼90%_ SBD2_369 / 451_ Alexa555647 samples according to ALEX sorting, recorded the apo-(ligand-free) and holo-(100 µM L-glutamine) conformation.

**Figure S5:**
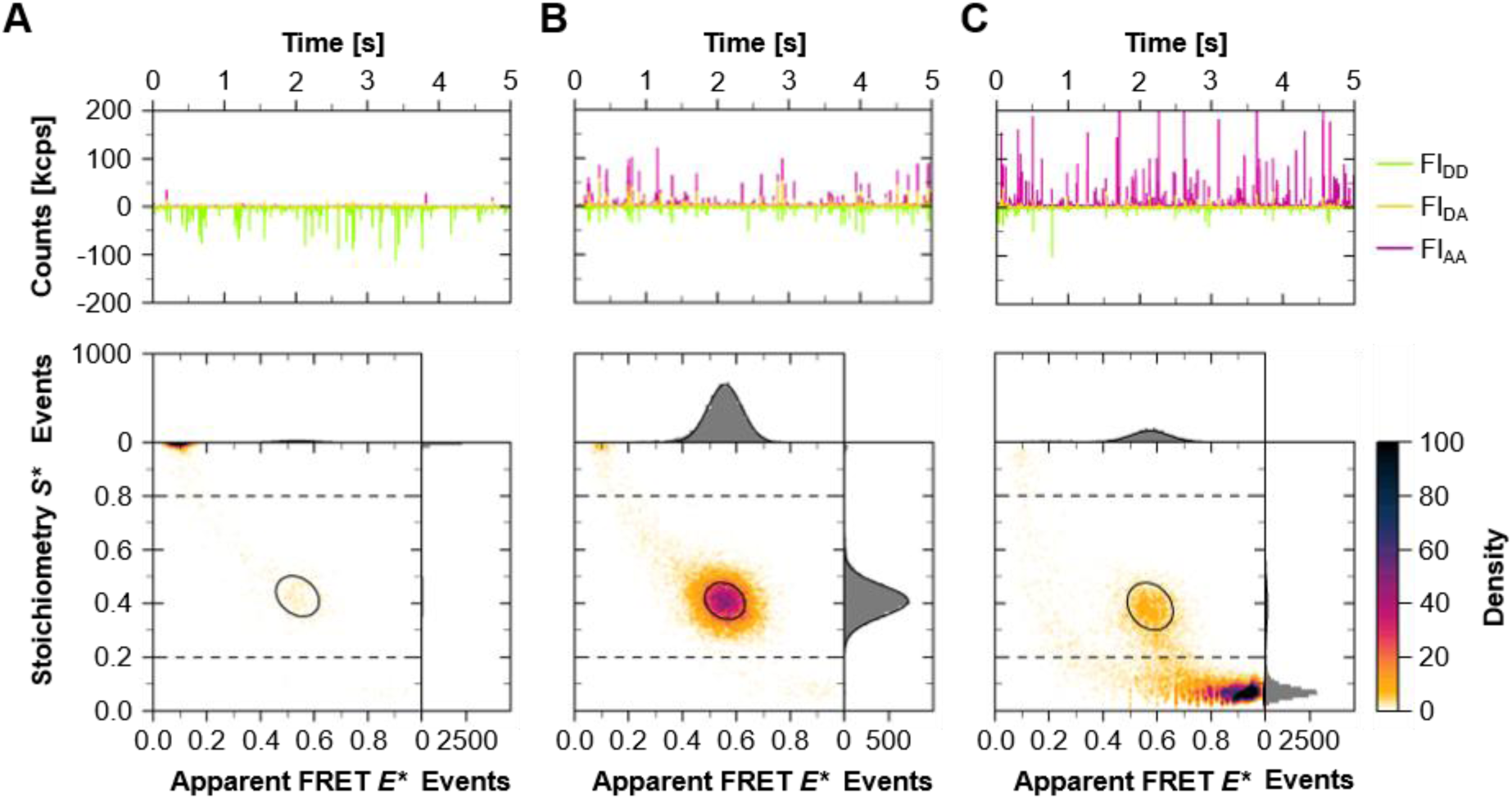
Data of SBD2_369 / 451_ with Alexa 555/647 as shown in. **Figure 2 but obtained with a MonoQ 5/50 GL**. A – C: Fluorescent time traces at 100 pM concentration and associated *E*S\**-histograms of the individual DD, DA, and DA populations, measured in the apo-(ligand-free) conformation.

**Figure S6:**
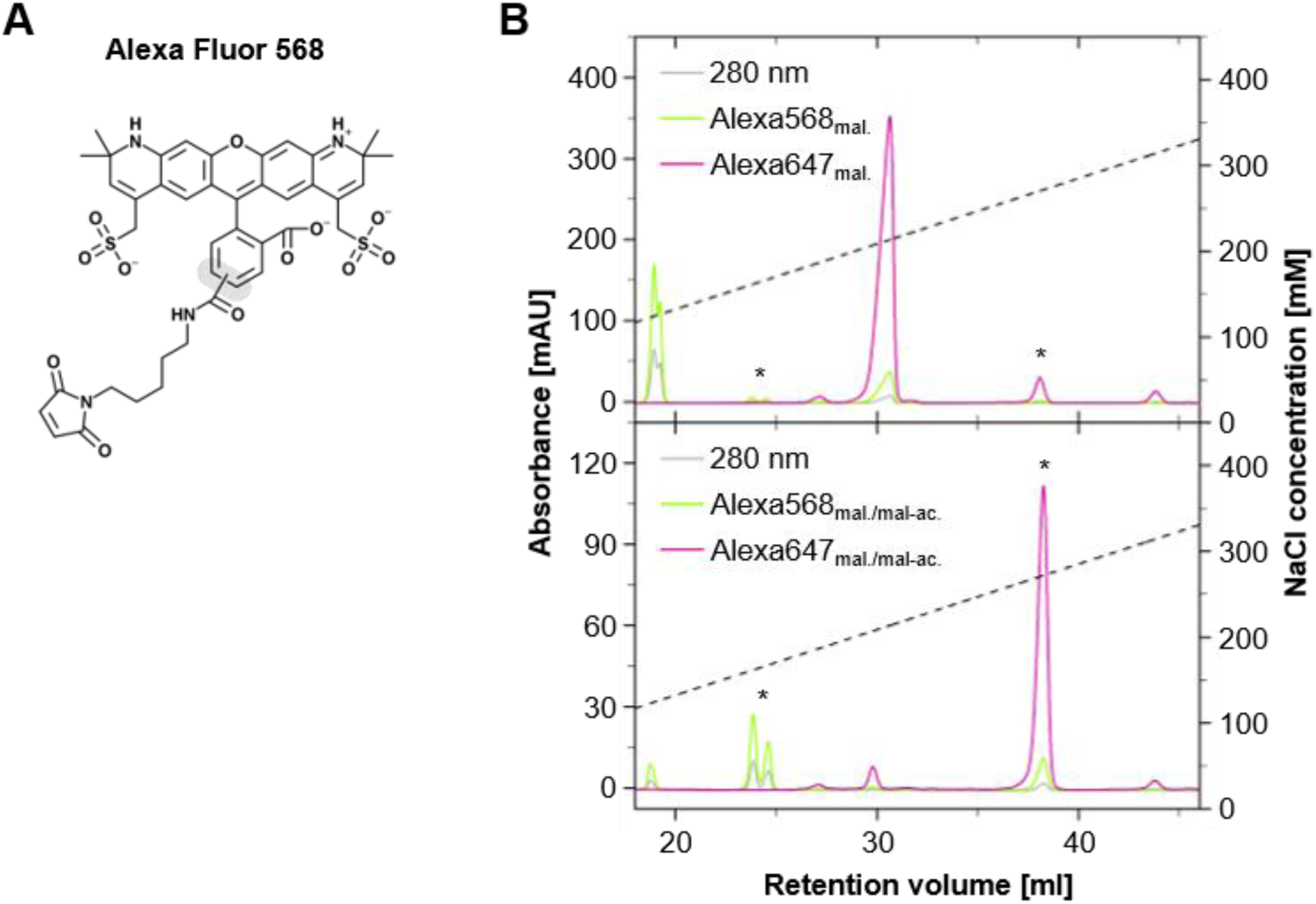
AIEX characterization (Capto HiRes Q 5/50) of the Alexa568/647 dye pair. A: Chemical structure of the isomeric Alexa568 maleimide (Thermo Fisher Scientific, [1]). B: Elution patterns during gradual elution (pH 7.5, 7.6 mM NaCl/ ml) of maleimide (mal.) and maleamic acid (*mal-ac.) versions of the individual fluorophores present in non-hydrolyzed (top panel) and partially hydrolyzed (lower panel) dye mixtures, based on absorbances measured at 280 nm (control wavelength), 575 nm (Alexa568), and 650 nm (Alexa647).

**Figure S7:**
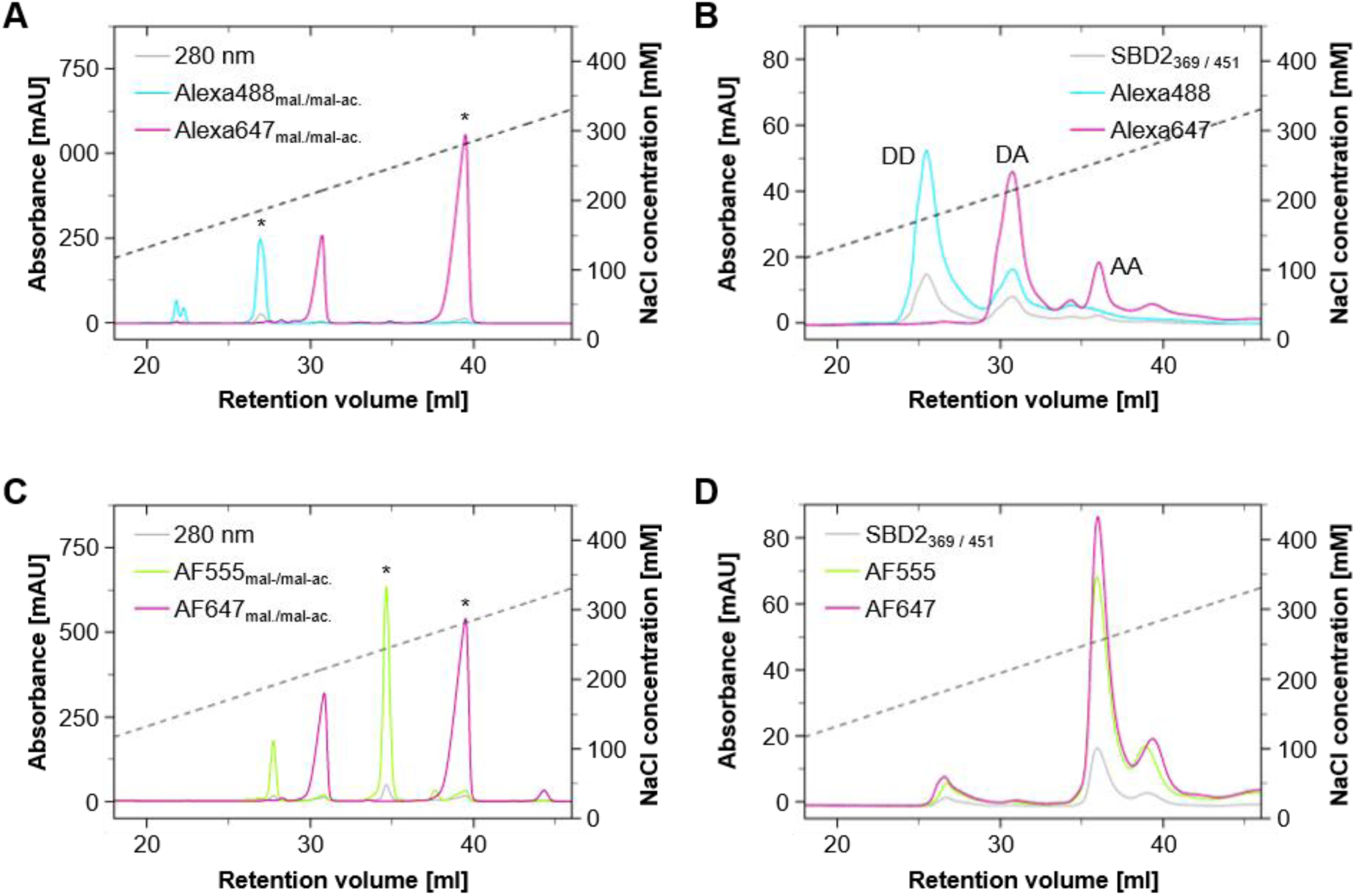
**Fluorophore profiles on AIEX and effects for AIEX separation on proteins**. A/C: Absorbance profiles of the ion dependent AIEX elution (pH 7.5, 7.6 mM NaCl/ ml) of the partially hydrolyzed (maleimide/mal. and *maleamic acid/mal-ac.) fluorophore pairs Alexa488/647 (439/650 nm), and AF555/647 (556/647 nm) using 14 nmol per fluorophore. B/D: Ion concentration dependent elution profiles (pH 8.5, 7.6 mM NaCl/ ml.) for SBD2_269 / 451_ (280 nm) labeled with the different fluorophore pairs.

**Figure S8:**
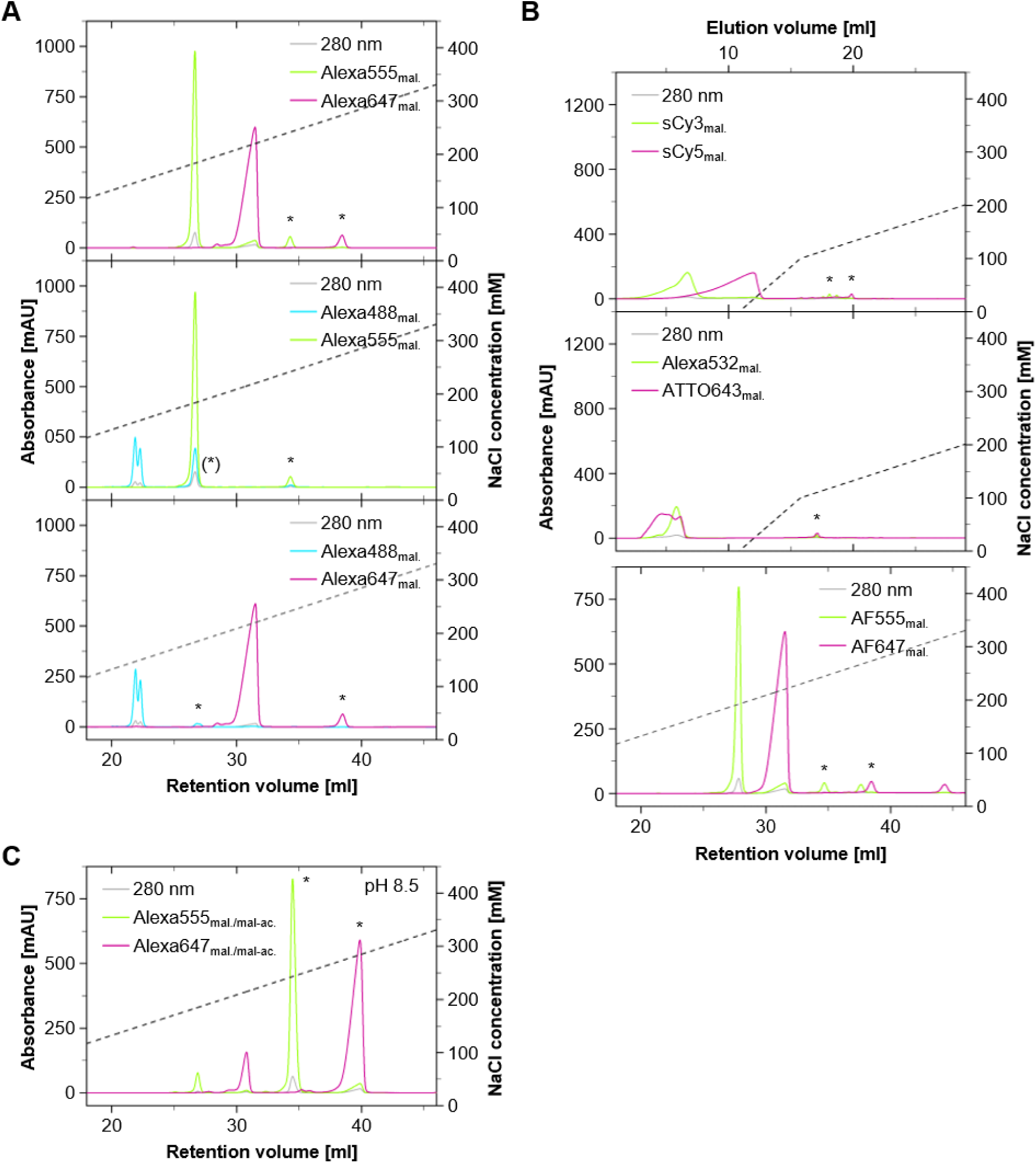
AIEX characterization of anionic fluorophore-maleimide pairs on the Capto HiRes Q 5/50. Non-hydrolyzed samples of 14 nmol contain minor amounts of maleamic acid (*mal-ac.) versions of each dye. Elution patterns during gradual elution (pH 7.5 or 8.5, 7.6 mM NaCl/ ml) are based on absorbances measured at 280 nm (control), 493 nm (Alexa488), 556 nm (Alexa555), 650 nm (Alexa647), 548 nm (sCy3), 646 nm (sCy5), 528 nm (Alexa532), 643 nm (ATTO643). A: Fluorophore-pairs that allow for the electrostatically dependent separation of differently labeled SBD2_369 / 451_ species. B: Fluorophore-pairs that are not suitable for a AIEX separation of labelled SBD2_369 / 451_ species. C: AIEX profile recorded at pH 8.5 of the partially hydrolyzed Alexa555/647 fluorophore pair.

**Figure S9:**
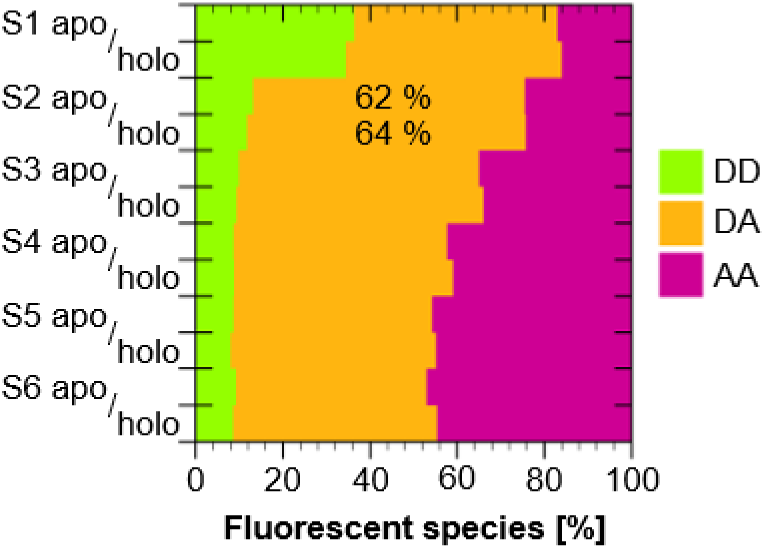
Quantification of fluorophore species fractions of SBD2_369 / 451_ with Alexa532/ATTO643. DD (including single D), DA, and AA (including single A) species contained in the different AIEX elution fractions S1-S6 according to ALEX sorting, recorded in the absence (apo) and presence of 100 µM L-glutamine (holo).

**Figure S10:**
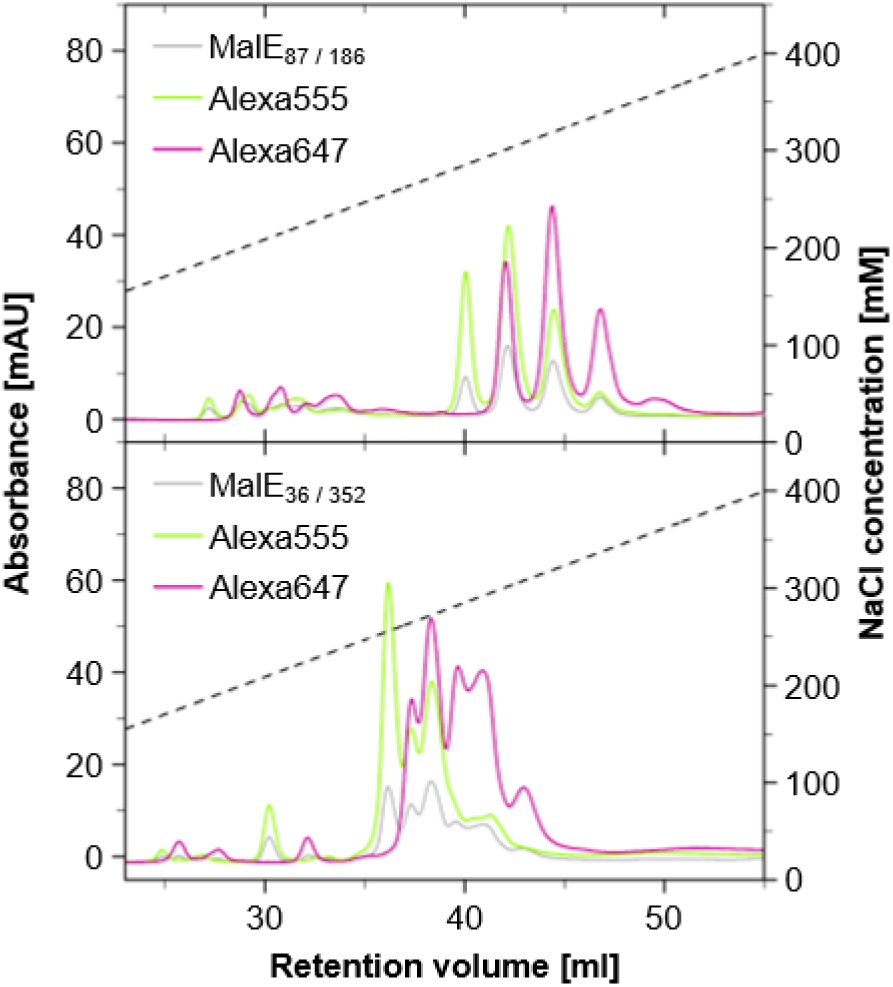
Profiles of AIEX purification of MalE_87 / 186_ and MalE_36 / 352_ with Alexa555/647 on the Capto HiRes Q 5/50. Elution patterns during gradual elution (pH 8.5, 7.6 mM NaCl/ ml, without initial NaCl supplementation) are based on absorbances measured at 280 nm (protein), 556 nm (Alexa555), and 650 nm (Alexa647).

**Figure S11:**
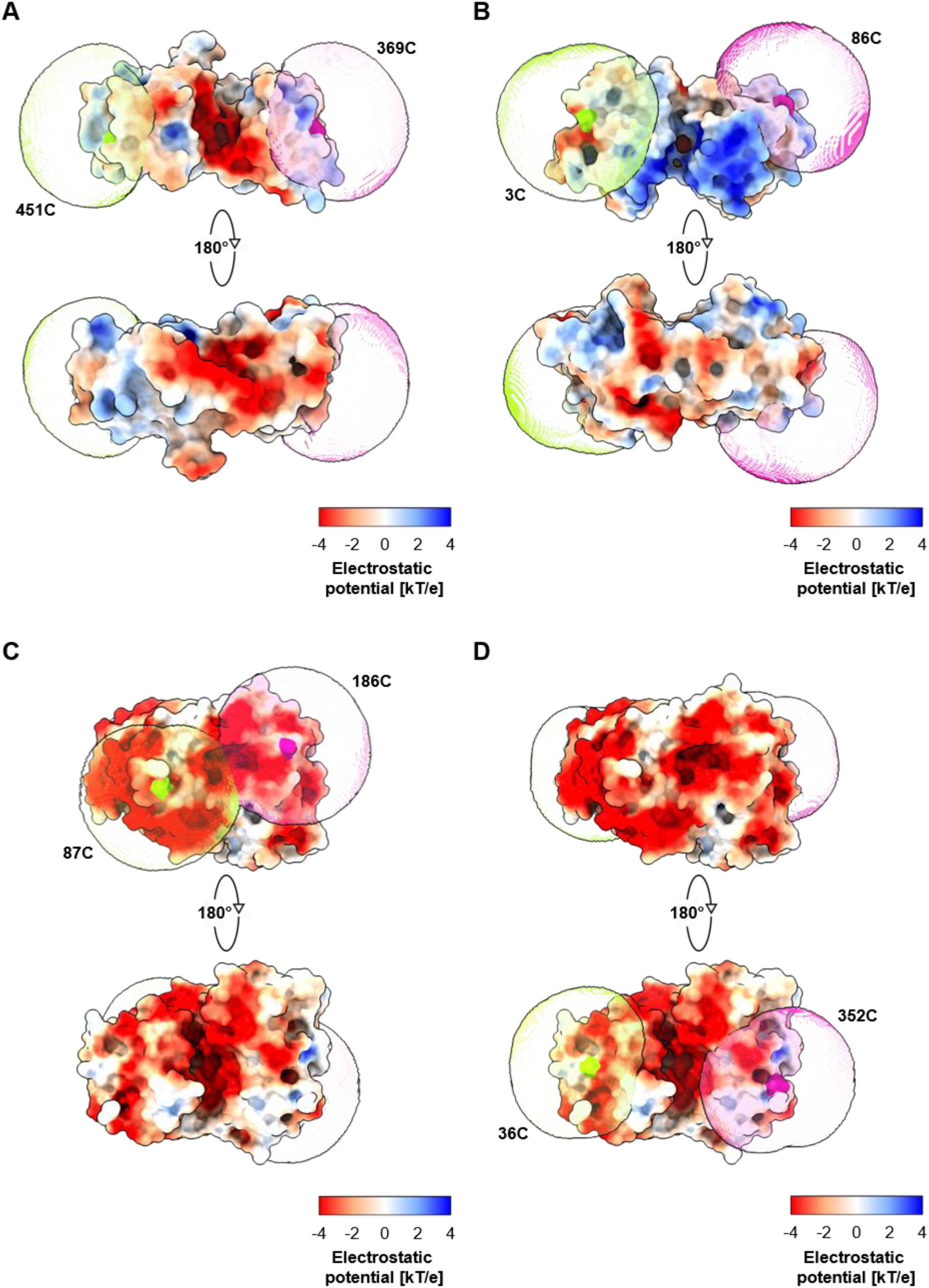
Surface potentials based on continuum electrostatic predictions of A: SBD2_369 / 451_ (PDB: 4KR5), B: PBP I76G_3 / 86_ (PDB: 1OIB), C: MalE_86 / 186_ (PDB: 1OMP), and D: MalE_36 / 352_ (PDB: 1OMP) at pH 8.5. The structural data was refined using the PDB2PQR software [2–7] in preparation for the continuum electrostatic calculations using the Adaptive Poisson Boltzmann Solver [8–10,7]. Fluorophore accessible volumes (AVs) are based on coarse-grained simulations via the FRET-restrained positioning and screening system [11] and are depicted for Alexa555 (green) and Alexa647 (red) fluorophores attached to the respective cysteine positions within the SBPs backbone.

**Figure S12:**
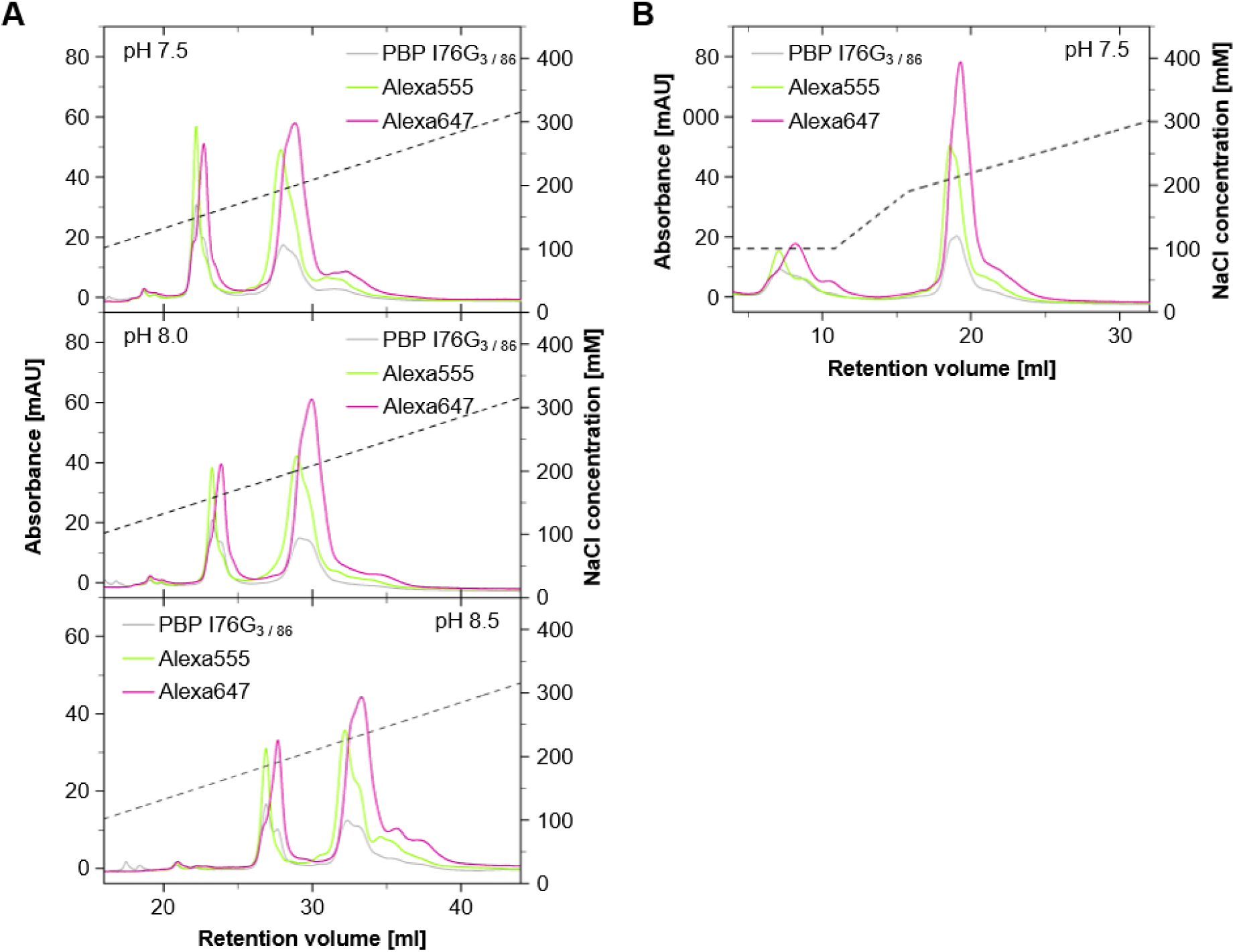
Evaluation of potential protein charge dependent influences on the AIEX purification with Capto HiRes Q 5/50 of stochastically labeled PBP I76G_3/ 86_ Alexa555/Alexa647 species. Elution patterns during gradual elution (7.6 mM NaCl/ ml) are based on absorbances measured at 280 nm (protein), 556 nm (Alexa555), and 650 nm (Alexa647). Significant amounts of singly-labelled (Alexa555 or Alexa647) PBP I76G_3/ 86_ are eluted earlier and independently of pH. A: The pH dependent shift of the PBP I76G_3/ 86_ Alexa555/647 elution profile according to the pH dependent negative protein surface charge potential. B: Effect of an initial supplementation with 100 mM NaCl on the elution pattern of the labelled PBP I76G_3 / 86_ at pH 7.5.

**Figure S13:**
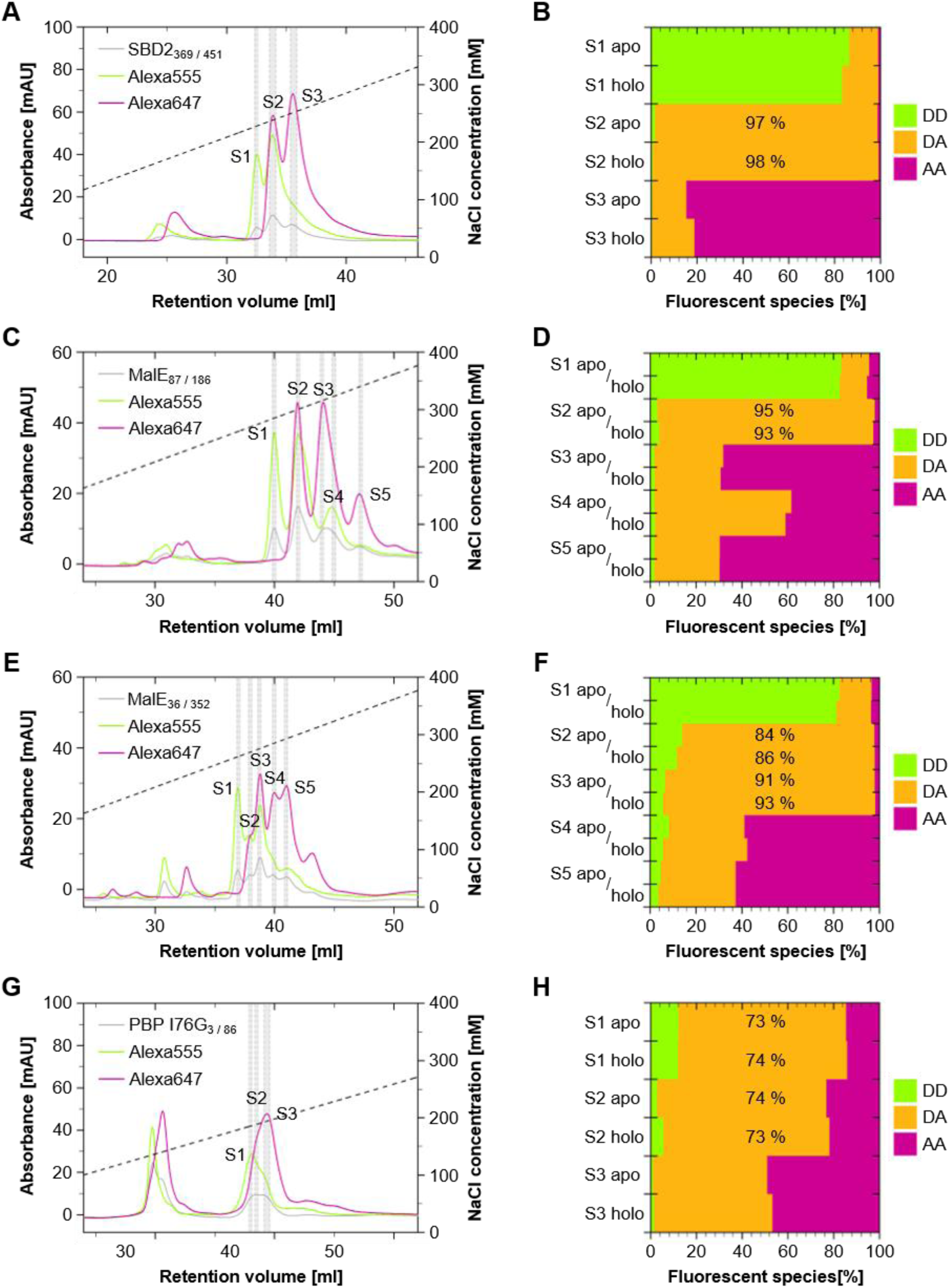
Comparative AIEX purification with a MonoQ 5/50 GL and characterization via ALEX. A, C, E: Absorbance profiles of the ion dependent separation (pH 8.5, 7.6 mM NaCl/ ml) of the different double labelled SBD2_369 / 451_ (S1-S3), MalE_87 / 186_ (S1-S5), MalE_36 / 352_ (S1–S5). G: Absorbance profile of the ion dependent elution (pH 8.5, 5.1 mM NaCl/ ml) of Alexa555 (556 nm)/647 (650 nm) labelled PBP I76G_3/ 86_ (280 nm) species (S1-S3). B, D, F, H: Relative population of labelled SBD2_369 / 451_, MalE_87 / 186_, MalE_36 / 352_, and PBP I76G_3 / 86_ species contained in the different AIEX elution fractions according to ALEX sorting, in both the apo-and holo-(SBD2: 100 µM L-glutamine; MalEs: 1mM D-maltose; PBP: 500 µM phosphate) conformation.

**Figure S14.**
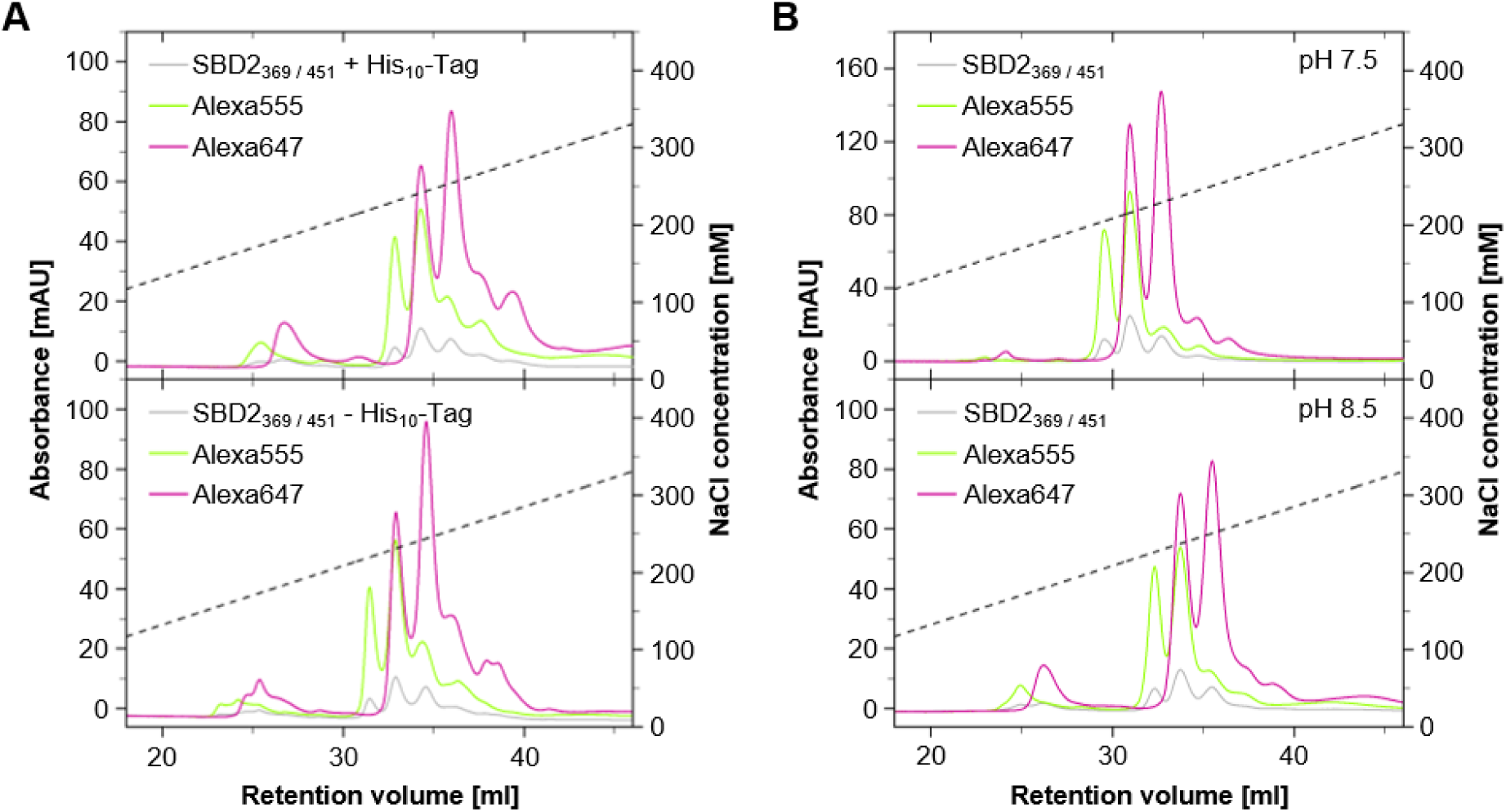
Evaluation of the influence of protein charge and purification tags (His_10_-Tag) on the Capto HiRes Q 5/50. Elution patterns during gradual elution (pH 8.5, 7.6 mM NaCl/ ml) of SBD2_369 / 451_ with Alexa555/647 species are based on absorbances measured at 280 nm (protein), 556 nm (Alexa555), and 650 nm (Alexa647). A: Influence of the N-terminal His_10_-Tag on the ion concentration dependent elution. B: The pH dependent shift of the elution profile on based on the pH dependent negative protein surface charge of the protein.

## Supplementary Note 1: Details on Material and Methods

### Expression and purification of recombinant substrate binding proteins MalE and SBD2

The expression and purification of maltose binding protein (MalE) variants MalE_T36C / S352C_ and MalE_D87C / A186C_ as well as GlnPQ substrate binding domain 2 (SBD2)_T369C / S451C_ generally followed already established and published protocols[12–14]. The T7*lac* controlled expression of MalE derivatives utilized the bacterial expression vector pET-20b(+) (Novagen) including a C-terminal His_6_-Tag fusion, for SBD2 a pBAD expression system was selected introducing a N-terminal His_10_-Tag extension. BL21(DE3)pLysS (Novagen) chemically competent cells were transformed with the respective vectors containing the coding sequences for recombinant MalE, or SBD2. Expression cultures were established in LB medium (Luria/ Miller) supplemented with carbenicillin (0.1 mg/ml) and chloramphenicol (0.05 mg/ml). A *lacUV5* controlled transcription of MalE was initially suppressed by the addition of 55 mM D-glucose. Protein-expression was induced by addition of 0.25 mM isopropyl β-D-1-thiogalactopyranoside (IPTG) for pET-20b(+), or 14 mM L-arabinose for the pBAD expression system. The cells were grown at 37°C for2 h, resuspended in 50 ml lysis buffer (50 mM Tris-HCl pH 7.4 for MalE/ pH 7.6 for SBD2, 1 M KCl, 10 % glycerol, 10 mM imidazole, 1 mM dithiothreitol), EDTA-free protease inhibitor cocktail (cOmplete, Roche), 0.2 mM phenylmethylsulfonyl fluoride (PMSF), and DNase I (500 ug/ml) were added and proteins were extracted by cell lysis using an ultrasonic homogenizer (Digital Sonifier 250, Branson Ultrasonics) equipped with a 5 mm diameter micro-tip probe (Branson Ultrasonics). The SBPs were purified by ultracentrifugation (208,400 × *g*, 1 h, 4 °C) and metal affinity chromatography (Ni^2+^-Sepharose 6 Fast Flow, Cytiva) and eluted in elution buffer (50 mM Tris-HCl pH 7.4 for MalE/ pH 7.6 for SBD2, 50 mM KCl, 10 % glycerol, 250 mM imidazole, 1 mM dithiothreitol). The eluates were dialyzed (10 KDa MWCO SnakeSkin, ThermoScientific) overnight at 4 °C twice,against 100 volumes of dialysis buffer (50 mM Tris-HCl 7.4 for MalE/ 7.6 for SBD2, 50 mM KCl, 1 mM dithiothreitol) and against 100 volumes of storage buffer (50 mM Tris-HCl pH 7.4 for MalE/ 7.6 for SBD2, 50 mM KCl, 50% glycerol, 1 mM dithiothreitol), Concentrations were determined by absorbance measurements (NanoPhotometer N60, IMPLEN) using the molar extinction coefficients 66350 l*mol^-1^*cm^-1^ for MalE derivatives (MW = 42.38 kDa), and 30370 l*mol^-1^*cm^-1^ for SBD2 (MW = 27.82 kDa).

### Expression and purification of recombinant phosphate binding protein (PBP)

For the expression and purification of phosphate binding protein (PBP) derivate PBP I76G_S3C / P86C_ the pET-20b(+) vector in combination with a BL21(DE3)pLysS host cell line was selected, and was carried out based on a modified protocol following the general order of the process described for MalE and SBD2 that has also been published before[15]. In short, adaptations include the cultivation in TB medium, the induction of over-expression with 0.5 mM IPTG and the harvest after 16 h. Additionally, the lysis buffer composition was adjusted (20 mM Hepes pH 7.5, 300 mM NaCl, 10 % glycerol, 10 mM imidazole, 1 mM dithiothreitol). During metal affinity chromatography, immobilized PBP was washed with wash buffer (20 mM Hepes pH 7.5, 300 mM NaCl, 10 % glycerol, 15 mM imidazole, 1 mM dithiothreitol) and the protein was eluted in PBP elution buffer (20 mM Hepes pH 7.5, 300 mM NaCl, 10 % glycerol, 500 mM imidazole, 1 mM dithiothreitol). The eluates were dialyzed (10 KDa MWCO SnakeSkin, ThermoScientific) overnight at 4 °C against 100 volumes of dialysis buffer (20 mM Tris-HCl 8.0, 100 mM NaCl, 1 mM dithiothreitol). Concentration calculations were based on a molar extinction coefficient of 61880 l*mol^-1^*cm^-1^ and a 35.45 kDa molecular weight.

### TEV protease mediated removal of poly-histidine tag from SBD2_369 / 451_ Alexa555/647

Site specific removal of the N-terminal His_10_-Tag from Alexa555/Alexa647 labelled SBD2_369 /451_ was facilitated using a TEV protease reaction. For the reaction a target protein to TEV protease ratio of 1:100 was chosen and the reaction was carried out in 50 mM Tris-HCl pH 7.6, 150 mM NaCl containing 0.5 mM EDTA. Digestions were started with a 2-hour incubation at 30 °C followed by a dialysis over night at 4°C against 1000 volumes of buffer containing 50 mM Tris-HCl pH 7.6 and 150 mM NaCl. Samples were concentrated and purified by AIEX. Restriction efficiency was evaluated using SDS gel electrophoresis with Coomassie staining.

### µsALEX measurements

Samples for solution based µsALEX confocal smFRET measurements followed a uniform workflow including data acquisition, data processing and data evaluation. Labelled and purified SBP species were measured on BSA passivated coverslips (1 mg/ml BSA in SBP specific buffer for 1 minute) in 100 µl droplets of SBP specific buffer (SBD2: 50 mM Tris-HCl pH 7.6, 150 mM NaCl/ MalE: 50 mM Tris-HCl pH 7.4, 50 mM KCl/ PBP: 20 mM Tris-HCl pH 8.0, 100 mM NaCl) and a final concentration between 50-150 pM. All measurements were performed on a homebuilt µsALEX setup[16]. For the excitation of the donor and acceptor fluorophores a 532 nm diode laser (OBIS 532-100-LS, Coherent) operated at 60 μW and a 637 nm diode laser (OBIS 637-140LX, Coherent) adjusted to 25 μW were used, respectively. The lasers were alternated at 20 kHz frequency, combined (T600lpxr, Chroma), and coupled into a single-mode fiber (P3-488PM-FC-2, Thorlabs). The incident laser light was guided to the confocal microscope via a dual edge beam splitter (ZT532/640rpc, Chroma) and focused to the excitation spot by a water immersion objective (UPlanSAPO 60x/1.2w, Olympus). The fluorescent sample emission was spatially filtered with a 50 μm pinhole, spectrally split (ZT640rdc, Chroma) into a donor and an acceptor channel, and individually filtered for (donor channel: BrightLine FF01-582/75-25, Semrock; acceptor channel: ET700/75m, Chroma) APD detection (SPCM-AQRH-14/ SPCM-AQR-H-34, Excelitas). Data analysis was performed using an in house developed software package[12].

### Fluorescence spectroscopy

Fluorescence emission spectra were recorded in SBP specific buffers (SBD2: 50 mM Tris-HCl pH 7.6, 150 mM NaCl/ MalE: 50 mM Tris-HCl pH 7.4, 50 mM KCl/ PBP: 20 mM Tris-HCl pH 8.0, 100 mM NaCl) on a fluorescence spectrometer (LS 55, Perkin Elmer) with excitation/emission slit width of 5/5 nm.

## Footnotes

1 SBD2 (apo: 4KR5; holo: 4KQP), MalE (apo: 1OMP; holo: 1ANF) and PBP (apo: 1OIB; holo: 1IXH)

## REFERENCES

1. Miller, H., Zhou, Z., Shepherd, J., Wollman, A.J.M., and Leake, M.C. (2017) Single-molecule techniques in biophysics: a review of the progress in methods and applications. Rep. Prog. Phys., 81 (2), 024601.

2. Deniz, A.A., Mukhopadhyay, S., and Lemke, E.A. (2007) Single-molecule biophysics: at the interface of biology, physics and chemistry. J R Soc Interface, 5 (18), 15–45.

3. Lerner, E., Cordes, T., Ingargiola, A., Alhadid, Y., Chung, S., Michalet, X., and Weiss, S. (2018) Toward dynamic structural biology: Two decades of single-molecule Förster resonance energy transfer. Science, 359 (6373), eaan1133.

4. Kapanidis, A.N., and Strick, T. (2009) Biology, one molecule at a time. Trends in Biochemical Sciences, 34 (5), 234–243.

5. Walter, N.G., Huang, C.-Y., Manzo, A.J., and Sobhy, M.A. (2008) Do-it-yourself guide: how to use the modern single-molecule toolkit. Nat Methods, 5 (6), 475–489.

6. Bernal, J.D., and Crowfoot, D. (1934) X-Ray Photographs of Crystalline Pepsin. Nature, 133 (3369), 794–795.

7. Zewail, A.H. (2004) Diffraction, crystallography and microscopy beyond three dimensions: structural dynamics in space and time. Philos Trans A Math Phys Eng Sci, 363 (1827), 315–329.

8. Bharat, T.A.M., Russo, C.J., Löwe, J., Passmore, L.A., and Scheres, S.H.W. (2015) Advances in Single-Particle Electron Cryomicroscopy Structure Determination applied to Sub-tomogram Averaging. Structure, 23 (9), 1743–1753.

9. Cheng, Y. (2015) Single-Particle Cryo-EM at Crystallographic Resolution. Cell, 161 (3), 450–457.

10. Dyson, H.J., and Wright, P.E. (2004) Unfolded Proteins and Protein Folding Studied by NMR. Chem. Rev., 104 (8), 3607–3622.

11. Jensen, M.R., Ruigrok, R.W., and Blackledge, M. (2013) Describing intrinsically disordered proteins at atomic resolution by NMR. Current Opinion in Structural Biology, 23 (3), 426–435.

12. Jensen, M.R., Zweckstetter, M., Huang, J., and Blackledge, M. (2014) Exploring Free-Energy Landscapes of Intrinsically Disordered Proteins at Atomic Resolution Using NMR Spectroscopy. Chem. Rev., 114 (13), 6632–6660.

13. Kaplan, M., Pinto, C., Houben, K., and Baldus, M. (2016) Nuclear magnetic resonance (NMR) applied to membrane–protein complexes. Quarterly Reviews of Biophysics, 49, e15.

14. Ha, T., Enderle, T., Ogletree, D.F., Chemla, D.S., Selvin, P.R., and Weiss, S. (1996) Probing the interaction between two single molecules: fluorescence resonance energy transfer between a single donor and a single acceptor. Proceedings of the National Academy of Sciences, 93 (13), 6264–6268.

15. Forster, Th. (1946) Energiewanderung und Fluoreszenz. Naturwissenschaften, 33 (6), 166–175.

16. Förster, Th. (1948) Zwischenmolekulare Energiewanderung und Fluoreszenz. Annalen der Physik, 437 (1–2), 55–75.

17. Roy, R., Hohng, S., and Ha, T. (2008) A practical guide to single-molecule FRET. Nat Methods, 5 (6), 507–516.

18. Stryer, L., and Haugland, R.P. (1967) Energy transfer: a spectroscopic ruler. Proceedings of the National Academy of Sciences, 58 (2), 719–726.

19. Lee, N.K., Kapanidis, A.N., Wang, Y., Michalet, X., Mukhopadhyay, J., Ebright, R.H., and Weiss, S. (2005) Accurate FRET Measurements within Single Diffusing Biomolecules Using Alternating-Laser Excitation. Biophysical Journal, 88 (4), 2939–2953.

20. Gilardi, Gianfranco., Zhou, L.Qing., Hibbert, Linda., and Cass, A.E.G. (1994) Engineering the Maltose Binding Protein for Reagentless Fluorescence Sensing. Anal. Chem., 66 (21), 3840–3847.

21. Brune, M., Hunter, J.L., Corrie, J.E.T., and Webb, M.R. (1994) Direct, Real-Time Measurement of Rapid Inorganic Phosphate Release Using a Novel Fluorescent Probe and Its Application to Actomyosin Subfragment 1 ATPase. Biochemistry, 33 (27), 8262–8271.

22. Marvin, J.S., Corcoran, E.E., Hattangadi, N.A., Zhang, J.V., Gere, S.A., and Hellinga, H.W. (1997) The rational design of allosteric interactions in a monomeric protein and its applications to the construction of biosensors. Proceedings of the National Academy of Sciences, 94 (9), 4366–4371.

23. Zhou, R., Kunzelmann, S., Webb, M.R., and Ha, T. (2011) Detecting Intramolecular Conformational Dynamics of Single Molecules in Short Distance Range with Subnanometer Sensitivity. Nano Lett., 11 (12), 5482–5488.

24. Marvin, J.S., and Hellinga, H.W. (1998) Engineering Biosensors by Introducing Fluorescent Allosteric Signal Transducers: Construction of a Novel Glucose Sensor. J. Am. Chem. Soc., 120 (1), 7–11.

25. Fitzgerald, G.A., Terry, D.S., Warren, A.L., Quick, M., Javitch, J.A., and Blanchard, S.C. (2019) Quantifying secondary transport at single-molecule resolution. Nature, 575 (7783), 528–534.

26. Lerner, E., Barth, A., Hendrix, J., Ambrose, B., Birkedal, V., Blanchard, S.C., Börner, R., Sung Chung, H., Cordes, T., Craggs, T.D., Deniz, A.A., Diao, J., Fei, J., Gonzalez, R.L., Gopich, I.V., Ha, T., Hanke, C.A., Haran, G., Hatzakis, N.S., Hohng, S., Hong, S.-C., Hugel, T., Ingargiola, A., Joo, C., Kapanidis, A.N., Kim, H.D., Laurence, T., Lee, N.K., Lee, T.-H., Lemke, E.A., Margeat, E., Michaelis, J., Michalet, X., Myong, S., Nettels, D., Peulen, T.-O., Ploetz, E., Razvag, Y., Robb, N.C., Schuler, B., Soleimaninejad, H., Tang, C., Vafabakhsh, R., Lamb, D.C., Seidel, C.A., and Weiss, S. (2021) FRET-based dynamic structural biology: Challenges, perspectives and an appeal for open-science practices. eLife, 10, e60416.

27. Ha, T., Kaiser, C., Myong, S., Wu, B., and Xiao, J. (2022) Next generation single-molecule techniques: Imaging, labeling, and manipulation in vitro and in cellulo. Mol Cell, 82 (2), 304–314.

28. Kahlscheuer, M.L., Widom, J., and Walter, N.G. (2015) Single-Molecule Pull-down FRET (SiMPull-FRET) to dissect the mechanisms of biomolecular machines. Methods Enzymol, 558, 539–570.

29. Selvin, P.R. (2000) The renaissance of fluorescence resonance energy transfer. Nat Struct Mol Biol, 7 (9), 730–734.

30. Sakon, J.J., and Weninger, K.R. (2010) Detecting the conformation of individual proteins in live cells. Nat Methods, 7 (3), 203–205.

31. König, I., Zarrine-Afsar, A., Aznauryan, M., Soranno, A., Wunderlich, B., Dingfelder, F., Stüber, J.C., Plückthun, A., Nettels, D., and Schuler, B. (2015) Single-molecule spectroscopy of protein conformational dynamics in live eukaryotic cells. Nat Methods, 12 (8), 773–779.

32. Sustarsic, M., and Kapanidis, A.N. (2015) Taking the ruler to the jungle: single-molecule FRET for understanding biomolecular structure and dynamics in live cells. Current Opinion in Structural Biology, 34, 52–59.

33. Anandamurugan, A., Eidloth, A., Frank, V., Wortmann, P., Schrangl, L., Lan, C., Schütz, G.J., and Hugel, T. (2025) Single-molecule FRET and tracking of transfected biomolecules in living cells. Biophysical Journal, 0 (0).

34. Lu, H.P. (2005) Probing Single-Molecule Protein Conformational Dynamics. Acc. Chem. Res., 38 (7), 557–565.

35. Henzler-Wildman, K.A., Thai, V., Lei, M., Ott, M., Wolf-Watz, M., Fenn, T., Pozharski, E., Wilson, M.A., Petsko, G.A., Karplus, M., Hübner, C.G., and Kern, D. (2007) Intrinsic motions along an enzymatic reaction trajectory. Nature, 450 (7171), 838–844.

36. Sasmal, D.K., Pulido, L.E., Kasal, S., and Huang, J. (2016) Single-molecule fluorescence resonance energy transfer in molecular biology. Nanoscale, 8 (48), 19928–19944.

37. Scheerer, D., Levy, D., Casier, R., Riven, I., Mazal, H., and Haran, G. (2025) Interplay between conformational dynamics and substrate binding regulates enzymatic activity: a single-molecule FRET study. Chemical Science, 16 (7), 3066–3077.

38. Zhao, R., and Rueda, D. (2009) RNA folding dynamics by single-molecule fluorescence resonance energy transfer. Methods, 49 (2), 112–117.

39. Christian, T.D., Romano, L.J., and Rueda, D. (2009) Single-molecule measurements of synthesis by DNA polymerase with base-pair resolution. Proceedings of the National Academy of Sciences, 106 (50), 21109–21114.

40. Preus, S., and Wilhelmsson, L.M. (2012) Advances in Quantitative FRET-Based Methods for Studying Nucleic Acids. ChemBioChem, 13 (14), 1990–2001.

41. Husada, F., Bountra, K., Tassis, K., de Boer, M., Romano, M., Rebuffat, S., Beis, K., and Cordes, T. (2018) Conformational dynamics of the ABC transporter McjD seen by single-molecule FRET. EMBO J, 37 (21), EMBJ2018100056.

42. Bartels, K., Lasitza-Male, T., Hofmann, H., and Löw, C. (2021) Single-Molecule FRET of Membrane Transport Proteins. ChemBioChem, 22 (17), 2657–2671.

43. Börsch, M., and Duncan, T.M. (2013) Spotlighting motors and controls of single FoF1-ATP synthase. Biochem Soc Trans, 41 (5), 1219–1226.

44. Schuler, B., and Eaton, W.A. (2008) Protein folding studied by single-molecule FRET. Current Opinion in Structural Biology, 18 (1), 16–26.

45. Borgia, A., Williams, P.M., and Clarke, J. (2008) Single-Molecule Studies of Protein Folding. Annu. Rev. Biochem., 77 (1), 101–125.

46. Gambin, Y., and Deniz, A.A. (2010) Multicolor single-molecule FRET to explore protein folding and binding. Mol Biosyst, 6 (9), 1540–1547.

47. Schuler, B., and Hofmann, H. (2013) Single-molecule spectroscopy of protein folding dynamics—expanding scope and timescales. Current Opinion in Structural Biology, 23 (1), 36–47.

48. Baldwin, A.D., and Kiick, K.L. (2011) Tunable Degradation of Maleimide–Thiol Adducts in Reducing Environments. Bioconjugate Chem., 22 (10), 1946–1953.

49. Fontaine, S.D., Reid, R., Robinson, L., Ashley, G.W., and Santi, D.V. (2015) Long-Term Stabilization of Maleimide–Thiol Conjugates. Bioconjugate Chem., 26 (1), 145–152.

50. Ravasco, J.M.J.M., Faustino, H., Trindade, A., and Gois, P.M.P. (2019) Bioconjugation with Maleimides: A Useful Tool for Chemical Biology. Chemistry – A European Journal, 25 (1), 43–59.

51. Hiscocks, H.G., Pascali, G., and Ung, A.T. (2023) Evaluation of Substituted N-Aryl Maleimide and Acrylamides for Bioconjugation. AppliedChem, 3 (2), 256–278.

52. Tyagi, S., and Lemke, E.A. (2015) Single-molecule FRET and crosslinking studies in structural biology enabled by noncanonical amino acids. Current Opinion in Structural Biology, 32, 66–73.

53. Popp, M.W.-L. (2015) Site-Specific Labeling of Proteins via Sortase: Protocols for the Molecular Biologist, in Site-Specific Protein Labeling: Methods and Protocols (eds.Gautier, A., and Hinner, M.J.), Springer, New York, NY, pp. 185–198.

54. Eggeling, C., Berger, S., Brand, L., Fries, J.R., Schaffer, J., Volkmer, A., and Seidel, C.A.M. (2001) Data registration and selective single-molecule analysis using multi-parameter fluorescence detection. Journal of Biotechnology, 86 (3), 163–180.

55. Widengren, J., Kudryavtsev, V., Antonik, M., Berger, S., Gerken, M., and Seidel, C.A.M. (2006) Single-Molecule Detection and Identification of Multiple Species by Multiparameter Fluorescence Detection. Anal. Chem., 78 (6), 2039–2050.

56. Müller, B.K., Zaychikov, E., Bräuchle, C., and Lamb, D.C. (2005) Pulsed Interleaved Excitation. Biophysical Journal, 89 (5), 3508–3522.

57. Kapanidis, A.N., Lee, N.K., Laurence, T.A., Doose, S., Margeat, E., and Weiss, S. (2004) Fluorescence-aided molecule sorting: Analysis of structure and interactions by alternating-laser excitation of single molecules. Proceedings of the National Academy of Sciences, 101 (24), 8936–8941.

58. Lerner, E., Hilzenrat, G., Amir, D., Tauber, E., Garini, Y., and Haas, E. (2013) Preparation of homogeneous samples of double-labelled protein suitable for single-molecule FRET measurements. Anal Bioanal Chem, 405 (18), 5983–5991.

59. Zosel, F., Holla, A., and Schuler, B. (2022) Labeling of Proteins for Single-Molecule Fluorescence Spectroscopy. Methods Mol Biol, 2376, 207–233.

60. Kapanidis, A.N., and Weiss, S. (2002) Fluorescent probes and bioconjugation chemistries for single-molecule fluorescence analysis of biomolecules. The Journal of Chemical Physics, 117 (24), 10953–10964.

61. Ratner, V., Kahana, E., Eichler, M., and Haas, E. (2002) A General Strategy for Site-Specific Double Labeling of Globular Proteins for Kinetic FRET Studies. Bioconjugate Chem., 13 (5), 1163–1170.

62. Kim, Y., Ho, S.O., Gassman, N.R., Korlann, Y., Landorf, E.V., Collart, F.R., and Weiss, S. (2008) Efficient Site-Specific Labeling of Proteins via Cysteines. Bioconjugate Chem., 19 (3), 786–791.

63. Site-Specific Labeling of Proteins for Single-Molecule FRET Measurements Using Genetically Encoded Ketone Functionalities | Springer Nature Experiments.

64. Gouridis, G., Schuurman-Wolters, G.K., Ploetz, E., Husada, F., Vietrov, R., de Boer, M., Cordes, T., and Poolman, B. (2015) Conformational dynamics in substrate-binding domains influences transport in the ABC importer GlnPQ. Nat Struct Mol Biol, 22 (1), 57–64.

65. de Boer, M., Gouridis, G., Vietrov, R., Begg, S.L., Schuurman-Wolters, G.K., Husada, F., Eleftheriadis, N., Poolman, B., McDevitt, C.A., and Cordes, T. (2019) Conformational and dynamic plasticity in substrate-binding proteins underlies selective transport in ABC importers. Elife, 8, e44652.

66. Gebhardt, C., Lehmann, M., Reif, M.M., Zacharias, M., Gemmecker, G., and Cordes, T. (2021) Molecular and Spectroscopic Characterization of Green and Red Cyanine Fluorophores from the Alexa Fluor and AF Series. ChemPhysChem, 22 (15), 1566–1583.

67. Agam, G., Gebhardt, C., Popara, M., Mächtel, R., Folz, J., Ambrose, B., Chamachi, N., Chung, S.Y., Craggs, T.D., de Boer, M., Grohmann, D., Ha, T., Hartmann, A., Hendrix, J., Hirschfeld, V., Hübner, C.G., Hugel, T., Kammerer, D., Kang, H.-S., Kapanidis, A.N., Krainer, G., Kramm, K., Lemke, E.A., Lerner, E., Margeat, E., Martens, K., Michaelis, J., Mitra, J., Moya Muñoz, G.G., Quast, R.B., Robb, N.C., Sattler, M., Schlierf, M., Schneider, J., Schröder, T., Sefer, A., Tan, P.S., Thurn, J., Tinnefeld, P., van Noort, J., Weiss, S., Wendler, N., Zijlstra, N., Barth, A., Seidel, C.A.M., Lamb, D.C., and Cordes, T. (2023) Reliability and accuracy of single-molecule FRET studies for characterization of structural dynamics and distances in proteins. Nat Methods, 20 (4), 523–535.

68. Gebhardt, C., Bawidamann, P., Spring, A.-K., Schenk, R., Schütze, K., Moya Muñoz, G.G., Wendler, N.D., Griffith, D.A., Lipfert, J., and Cordes, T. (2025) Labelizer: systematic selection of protein residues for covalent fluorophore labeling. Nat Commun, 16 (1), 4147.

69. Husada, F., Gouridis, G., Vietrov, R., Schuurman-Wolters, G.K., Ploetz, E., de Boer, M., Poolman, B., and Cordes, T. (2015) Watching conformational dynamics of ABC transporters with single-molecule tools. Biochem Soc Trans, 43 (5), 1041–1047.

70. Jazi, A.A., Ploetz, E., Arizki, M., Dhandayuthapani, B., Waclawska, I., Krämer, R., Ziegler, C., and Cordes, T. (2017) Caging and Photoactivation in Single-Molecule Förster Resonance Energy Transfer Experiments. Biochemistry, 56 (14), 2031–2041.

71. Dolinsky, T.J., Nielsen, J.E., McCammon, J.A., and Baker, N.A. (2004) PDB2PQR: an automated pipeline for the setup of Poisson–Boltzmann electrostatics calculations. Nucleic Acids Res, 32 (suppl_2), W665–W667.

72. Dolinsky, T.J., Czodrowski, P., Li, H., Nielsen, J.E., Jensen, J.H., Klebe, G., and Baker, N.A. (2007) PDB2PQR: expanding and upgrading automated preparation of biomolecular structures for molecular simulations. Nucleic Acids Research, 35 (Web Server), W522–W525.

73. Unni, S., Huang, Y., Hanson, R.M., Tobias, M., Krishnan, S., Li, W.W., Nielsen, J.E., and Baker, N.A. (2011) Web servers and services for electrostatics calculations with APBS and PDB2PQR. Journal of Computational Chemistry, 32 (7), 1488–1491.

74. Jurrus, E., Engel, D., Star, K., Monson, K., Brandi, J., Felberg, L.E., Brookes, D.H., Wilson, L., Chen, J., Liles, K., Chun, M., Li, P., Gohara, D.W., Dolinsky, T., Konecny, R., Koes, D.R., Nielsen, J.E., Head-Gordon, T., Geng, W., Krasny, R., Wei, G.-W., Holst, M.J., McCammon, J.A., and Baker, N.A. (2018) Improvements to the APBS biomolecular solvation software suite. Protein Science, 27 (1), 112–128.

75. Ploetz, E., Schuurman-Wolters, G.K., Zijlstra, N., Jager, A.W., Griffith, D.A., Guskov, A., Gouridis, G., Poolman, B., and Cordes, T. (2021) Structural and biophysical characterization of the tandem substrate-binding domains of the ABC importer GlnPQ. Open Biology, 11 (4), 200406.

76. Heitz, J.R., Anderson, C.D., and Anderson, B.M. (1968) Inactivation of yeast alcohol dehydrogenase by *N*-alkylmaleimides. Archives of Biochemistry and Biophysics, 127, 627–636.

77. Khan, M.N., and Khan, A.A. (1975) Kinetics and mechanism of hydrolysis of succinimide in highly alkaline medium. J. Org. Chem., 40 (12), 1793–1794.

78. Barradas, R.G., Fletcher, S., and Porter, J.D. (1976) The hydrolysis of maleimide in alkaline solution. Can. J. Chem., 54 (9), 1400–1404.

79. Yu, X.C., Joe, K., Zhang, Y., Adriano, A., Wang, Y., Gazzano-Santoro, H., Keck, R.G., Deperalta, G., and Ling, V. (2011) Accurate Determination of Succinimide Degradation Products Using High Fidelity Trypsin Digestion Peptide Map Analysis. Anal. Chem., 83 (15), 5912–5919.

80. Alexa Fluor^TM^ 532 C5 Maleimide | Invitrogen^TM^ | thermofisher.com.

81. Alexa Fluor^TM^ 488 C5 Maleimid | Invitrogen^TM^ | thermofisher.com.

82. sulfo-Cyanin3-Maleimid. Lumiprobe.

83. sulfo-Cyanine5 maleimide. Lumiprobe.

84. Vrljic, M., Strop, P., Ernst, J.A., Sutton, R.B., Chu, S., and Brunger, A.T. (2010) Molecular mechanism of the synaptotagmin–SNARE interaction in Ca2+-triggered vesicle fusion. Nat Struct Mol Biol, 17 (3), 325–331.

85. Alexa Fluor^TM^ 568 C5 Maleimide | Invitrogen^TM^ | thermofisher.com.

86. Hall, J.A., Ganesan, A.K., Chen, J., and Nikaido, H. (1997) Two Modes of Ligand Binding in Maltose-binding Protein ofEscherichia coli: FUNCTIONAL SIGNIFICANCE IN ACTIVE TRANSPORT *. Journal of Biological Chemistry, 272 (28), 17615–17622.

87. Kim, E., Lee, S., Jeon, A., Choi, J.M., Lee, H.-S., Hohng, S., and Kim, H.-S. (2013) A single-molecule dissection of ligand binding to a protein with intrinsic dynamics. Nat Chem Biol, 9 (5), 313–318.

88. Stubhan, S., V. Baptist, A., Körösy, C., Narducci, A., Muñoz, G.G.M., Wendler, N., Lak, A., Sztucki, M., Cordes, T., and Lipfert, J. (2025) Determination of absolute intramolecular distances in proteins using anomalous X-ray scattering interferometry. Nanoscale, 17 (6), 3322–3330.

89. Munoz, G.M., Luna, J., Con, P., Rohman, M.A., Lu, S., Peulen, T.O., and Cordes, T. (2025) Single-molecule FRET with a minimalistic 3D-printed setup and dyes in the blue-green spectral region. 2025.12.16.694555.

90. Kalinin, S., Peulen, T., Sindbert, S., Rothwell, P.J., Berger, S., Restle, T., Goody, R.S., Gohlke, H., and Seidel, C.A.M. (2012) A toolkit and benchmark study for FRET-restrained high-precision structural modeling. Nat Methods, 9 (12), 1218–1225.

91. Zhang, L., Isselstein, M., Köhler, J., Eleftheriadis, N., Huisjes, N.M., Guirao-Ortiz, M., Narducci, A., Smit, J.H., Stoffels, J., Harz, H., Leonhardt, H., Herrmann, A., and Cordes, T. (2022) Linker Molecules Convert Commercial Fluorophores into Tailored Functional Probes during Biolabelling. Angewandte Chemie, 134 (19), e202112959.

92. Roßmann, K., C. Akkaya, K., Poc, P., Charbonnier, C., Eichhorst, J., Gonschior, H., Valavalkar, A., Wendler, N., Cordes, T., Dietzek-Ivanšić, B., Jones, B., Lehmann, M., and Broichhagen, J. (2022) N-Methyl deuterated rhodamines for protein labelling in sensitive fluorescence microscopy. Chemical Science, 13 (29), 8605–8617.

93. Liphardt, B., Liphardt, B., and Lüttke, W. (1981) Laser dyes with intramolecular triplet quenching. Optics Communications, 38 (3), 207–210.

94. Schäfer, F.P., Zhang, F.-G., and Jethwa, J. (1982) Intramolecular TT-energy transfer in bifluorophoric laser dyes. Appl. Phys. B, 28 (1), 37–41.

95. Altman, R.B., Terry, D.S., Zhou, Z., Zheng, Q., Geggier, P., Kolster, R.A., Zhao, Y., Javitch, J.A., Warren, J.D., and Blanchard, S.C. (2012) Cyanine fluorophore derivatives with enhanced photostability. Nat Methods, 9 (1), 68–71.

96. Zheng, Q., Jockusch, S., Zhou, Z., Altman, R.B., Warren, J.D., Turro, N.J., and Blanchard, S.C. (2012) On the Mechanisms of Cyanine Fluorophore Photostabilization. J. Phys. Chem. Lett., 3 (16), 2200–2203.

97. van der Velde, J.H.M., Ploetz, E., Hiermaier, M., Oelerich, J., de Vries, J.W., Roelfes, G., and Cordes, T. (2013) Mechanism of Intramolecular Photostabilization in Self-Healing Cyanine Fluorophores. ChemPhysChem, 14 (18), 4084–4093.

98. van der Velde, J.H.M., Oelerich, J., Huang, J., Smit, J.H., Aminian Jazi, A., Galiani, S., Kolmakov, K., Gouridis, G., Eggeling, C., Herrmann, A., Roelfes, G., and Cordes, T. (2016) A simple and versatile design concept for fluorophore derivatives with intramolecular photostabilization. Nat Commun, 7 (1), 10144.

99. Isselstein, M., Zhang, L., Glembockyte, V., Brix, O., Cosa, G., Tinnefeld, P., and Cordes, T. (2020) Self-Healing Dyes—Keeping the Promise? J. Phys. Chem. Lett., 11 (11), 4462–4480.

100. Yang, Z., Li, L., Ling, J., Liu, T., Huang, X., Ying, Y., Zhao, Y., Zhao, Y., Lei, K., Chen, L., and Chen, Z. (2020) Cyclooctatetraene-conjugated cyanine mitochondrial probes minimize phototoxicity in fluorescence and nanoscopic imaging. Chemical Science, 11 (32), 8506–8516.

101. Zhang, Y., Yang, C., Peng, S., Ling, J., Chen, P., Ma, Y., Wang, W., Chen, Z., and Chen, C. (2023) General Strategy To Improve the Photon Budget of Thiol-Conjugated Cyanine Dyes. J. Am. Chem. Soc., 145 (7), 4187–4198.

102. Peter, M.F., Gebhardt, C., Mächtel, R., Muñoz, G.G.M., Glaenzer, J., Narducci, A., Thomas, G.H., Cordes, T., and Hagelueken, G. (2022) Cross-validation of distance measurements in proteins by PELDOR/DEER and single-molecule FRET. Nat Commun, 13 (1), 4396.

## SI References

1. Alexa Fluor^TM^ 568 C5 Maleimide 1 mg | Buy Online | Invitrogen^TM^ | thermofisher.com.

2. Holst, M., and Saied, F. (1993) Multigrid solution of the Poisson—Boltzmann equation. Journal of Computational Chemistry, 14 (1), 105–113.

3. Holst, M.J., and Saied, F. (1995) Numerical solution of the nonlinear Poisson–Boltzmann equation: Developing more robust and efficient methods. Journal of Computational Chemistry, 16 (3), 337–364.

4. Holst, M. (2001) Adaptive Numerical Treatment of Elliptic Systems on Manifolds. Advances in Computational Mathematics, 15 (1), 139–191.

5. Baker, N.A., Sept, D., Joseph, S., Holst, M.J., and McCammon, J.A. (2001) Electrostatics of nanosystems: Application to microtubules and the ribosome. Proceedings of the National Academy of Sciences, 98 (18), 10037–10041.

6. Bank, R.E., and Holst, M. (2003) A New Paradigm for Parallel Adaptive Meshing Algorithms. SIAM Rev., 45 (2), 291–323.

7. Jurrus, E., Engel, D., Star, K., Monson, K., Brandi, J., Felberg, L.E., Brookes, D.H., Wilson, L., Chen, J., Liles, K., Chun, M., Li, P., Gohara, D.W., Dolinsky, T., Konecny, R., Koes, D.R., Nielsen, J.E., Head-Gordon, T., Geng, W., Krasny, R., Wei, G.-W., Holst, M.J., McCammon, J.A., and Baker, N.A. (2018) Improvements to the APBS biomolecular solvation software suite. Protein Science, 27 (1), 112–128.

8. Dolinsky, T.J., Nielsen, J.E., McCammon, J.A., and Baker, N.A. (2004) PDB2PQR: an automated pipeline for the setup of Poisson–Boltzmann electrostatics calculations. Nucleic Acids Res, 32 (suppl_2), W665–W667.

9. Dolinsky, T.J., Czodrowski, P., Li, H., Nielsen, J.E., Jensen, J.H., Klebe, G., and Baker, N.A. (2007) PDB2PQR: expanding and upgrading automated preparation of biomolecular structures for molecular simulations. Nucleic Acids Research, 35 (Web Server), W522–W525.

10. Unni, S., Huang, Y., Hanson, R.M., Tobias, M., Krishnan, S., Li, W.W., Nielsen, J.E., and Baker, N.A. (2011) Web servers and services for electrostatics calculations with APBS and PDB2PQR. Journal of Computational Chemistry, 32 (7), 1488–1491.

11. Kalinin, S., Peulen, T., Sindbert, S., Rothwell, P.J., Berger, S., Restle, T., Goody, R.S., Gohlke, H., and Seidel, C.A.M. (2012) A toolkit and benchmark study for FRET-restrained high-precision structural modeling. Nat Methods, 9 (12), 1218–1225.

12. Gouridis, G., Schuurman-Wolters, G.K., Ploetz, E., Husada, F., Vietrov, R., de Boer, M., Cordes, T., and Poolman, B. (2015) Conformational dynamics in substrate-binding domains influences transport in the ABC importer GlnPQ. Nat Struct Mol Biol, 22 (1), 57–64.

13. de Boer, M., Gouridis, G., Vietrov, R., Begg, S.L., Schuurman-Wolters, G.K., Husada, F., Eleftheriadis, N., Poolman, B., McDevitt, C.A., and Cordes, T. (2019) Conformational and dynamic plasticity in substrate-binding proteins underlies selective transport in ABC importers. Elife, 8, e44652.

14. Peter, M.F., Gebhardt, C., Mächtel, R., Muñoz, G.G.M., Glaenzer, J., Narducci, A., Thomas, G.H., Cordes, T., and Hagelueken, G. (2022) Cross-validation of distance measurements in proteins by PELDOR/DEER and single-molecule FRET. Nat Commun, 13 (1), 4396.

15. Gebhardt, C., Bawidamann, P., Spring, A.-K., Schenk, R., Schütze, K., Moya Muñoz, G.G., Wendler, N.D., Griffith, D.A., Lipfert, J., and Cordes, T. (2025) Labelizer: systematic selection of protein residues for covalent fluorophore labeling. Nat Commun, 16 (1), 4147.

16. Hellenkamp, B., Schmid, S., Doroshenko, O., Opanasyuk, O., Kühnemuth, R., Rezaei Adariani, S., Ambrose, B., Aznauryan, M., Barth, A., Birkedal, V., Bowen, M.E., Chen, H., Cordes, T., Eilert, T., Fijen, C., Gebhardt, C., Götz, M., Gouridis, G., Gratton, E., Ha, T., Hao, P., Hanke, C.A., Hartmann, A., Hendrix, J., Hildebrandt, L.L., Hirschfeld, V., Hohlbein, J., Hua, B., Hübner, C.G., Kallis, E., Kapanidis, A.N., Kim, J.-Y., Krainer, G., Lamb, D.C., Lee, N.K., Lemke, E.A., Levesque, B., Levitus, M., McCann, J.J., Naredi-Rainer, N., Nettels, D., Ngo, T., Qiu, R., Robb, N.C., Röcker, C., Sanabria, H., Schlierf, M., Schröder, T., Schuler, B., Seidel, H., Streit, L., Thurn, J., Tinnefeld, P., Tyagi, S., Vandenberk, N., Vera, A.M., Weninger, K.R., Wünsch, B., Yanez-Orozco, I.S., Michaelis, J., Seidel, C.A.M., Craggs, T.D., and Hugel, T. (2018) Precision and accuracy of single-molecule FRET measurements—a multi-laboratory benchmark study. Nat Methods, 15 (9), 669–676.

